# Metabolic acids impact bone mineral maturation

**DOI:** 10.1101/2022.09.21.508894

**Authors:** Yang Li, Rui Li, David G. Reid, Joe T. Lunn, Karin H. Müller, Danielle Laurencin, Christian Bonhomme, E. Alex Ossa, Nico A.J.M. Sommerdijk, Melinda J Duer

## Abstract

Bone mineral has a complex 3D architecture that is essential to its mechanical properties. It is a complex calcium phosphate phase related to hydroxyapatite that also contains significant quantities of cell respiration metabolites, in particular: carbonate, citrate and lactate. An as-yet unanswered question is what, if any, role do these metabolites collectively play in determining the 3D architecture of bone mineral? Here we synthesize apatitic materials by transformation from precursor mineral phases containing citrate, lactate or carbonate so that the synthesis environment mimics the densely-packed ionic environment within which bone mineral forms in vivo, and so that we can understand the mineral factors that may direct bone mineral 3D architecture. We show that incorporating citrate and lactate leads to complex mineral architectures reminiscent of those in bone mineral, including curvature of the mineral crystals. Our results suggest that metabolic acids may assist the moulding of bone mineral to restricted spaces available for mineral in in vivo bone. We find that the incorporation of lactate creates a softer material and inhibits the transformation towards apatitic structures, which may help to explain why foetal bone – necessarily soft – contains considerable quantities of lactate. High levels of plasma citrate have been previously found to correlate with high bone mineral density. Here we find that citrate incorporation leads to mineral crystal curvature modelling that in in vivo bone mineral suggesting its importance in mineral morphology. We conclude that metabolic anions may play an important role in controlling bone mineral physicochemical properties and 3D architecture.

## Introduction

Bone mineral mechanical properties are essential to bone health. They depend on the mineral 3D architecture, the building blocks the mineral is made up of and how those building blocks are held together. Bone mineral forms in vivo within and around collagen fibrils in spaces generated by the specific arrangement of collagen molecules and the fibrils that form from them. Clearly then, the collagen matrix plays an important role in defining the spaces available for mineral crystals and thus makes a significant contribution to the mineral 3D architecture. A key, as-yet unanswered question is: is this the only factor determining the mineral architecture or are there components in the mineral itself which contribute to its shaping?

TEM studies show that the building blocks in bone are needle-shaped mineral crystals that are laterally aggregated and partially merged into ∼5 nm thick, curved platelets,^1,2^ the platelets being arranged in interlaced stacks of 2 – 4 platelets inside and around collagen fibrils.^1–5^ This complex mineral crystal organization is ultimately what leads to the overall mineral 3D architecture, but what holds the mineral crystals together in this specific arrangement is less clear. Clearly something drives the formation of the interfaces between the mineral needles to form platelets and the interfaces between those resulting platelets, but what is still an open question. Are the interfaces formed purely because the underlying collagen matrix forces the mineral crystals into these spatial relationships or is there a contribution from the chemistry of the mineral itself?

Bone mineral is a complex calcium phosphate phase related to hydroxyapatite (HAp). Importantly, it is not a purely inorganic material; it contains significant quantities of cell respiration metabolites: carbonate (6 - 8 wt%)^6^, citrate (2 – 5 wt%)^7–9^ and variable amounts of lactate.^7,8,10^ Bone mineral has been extensively modelled as nanocrystalline hydroxyapatite (HAp) and carbonate-substituted HAp.^11^ Nanocrystals of both HAp and carbonate-substituted HAp are flat, rectilinear lamellae. They do not resemble the curved, aggregated needle morphology of in vivo bone mineral. Thus, there is no obvious contribution from either HAp itself or from incorporation of carbonate to the morphology of bone mineral in the absence of a collagen matrix beyond the facility to form nanocrystals. Interestingly though, it has been shown that carbonate facilitates the formation of disordered, highly hydrated surface layers on HAp nanocrystals and it has been hypothesized that these layers may model interfaces between mineral platelets in in vivo bone [refs].

The roles of citrate and lactate in bone mineral structure have been much less studied than those of carbonate. In adult humans, bone mineral density correlates positively with the concentration of blood plasma citrate,^12^ whilst foetal bone, which is typically much softer than mature bone, contains considerable quantities of lactate.^13^ We have shown that the citrate chemical environment in bone is similar to that in octacalcium phosphate (OCP)-citrate by SSNMR spectroscopy.^14^ The citrate anions in OCP-citrate reside in disordered, highly hydrated layers, bridging between the OCP (OCP, Ca_16_(HPO_4_)_4_(PO_4_)_8_.10H_2_O) apatitic-like layers (Fig 1), suggesting that in bone, citrate anions may similarly reside at interfaces between apatitic mineral crystals. However, OCP-citrate is not a good model of the apatitic regions of bone mineral as it lacks the apatitic hydroxyl groups characteristic of bone mineral.^15,16^

**Fig 1:**
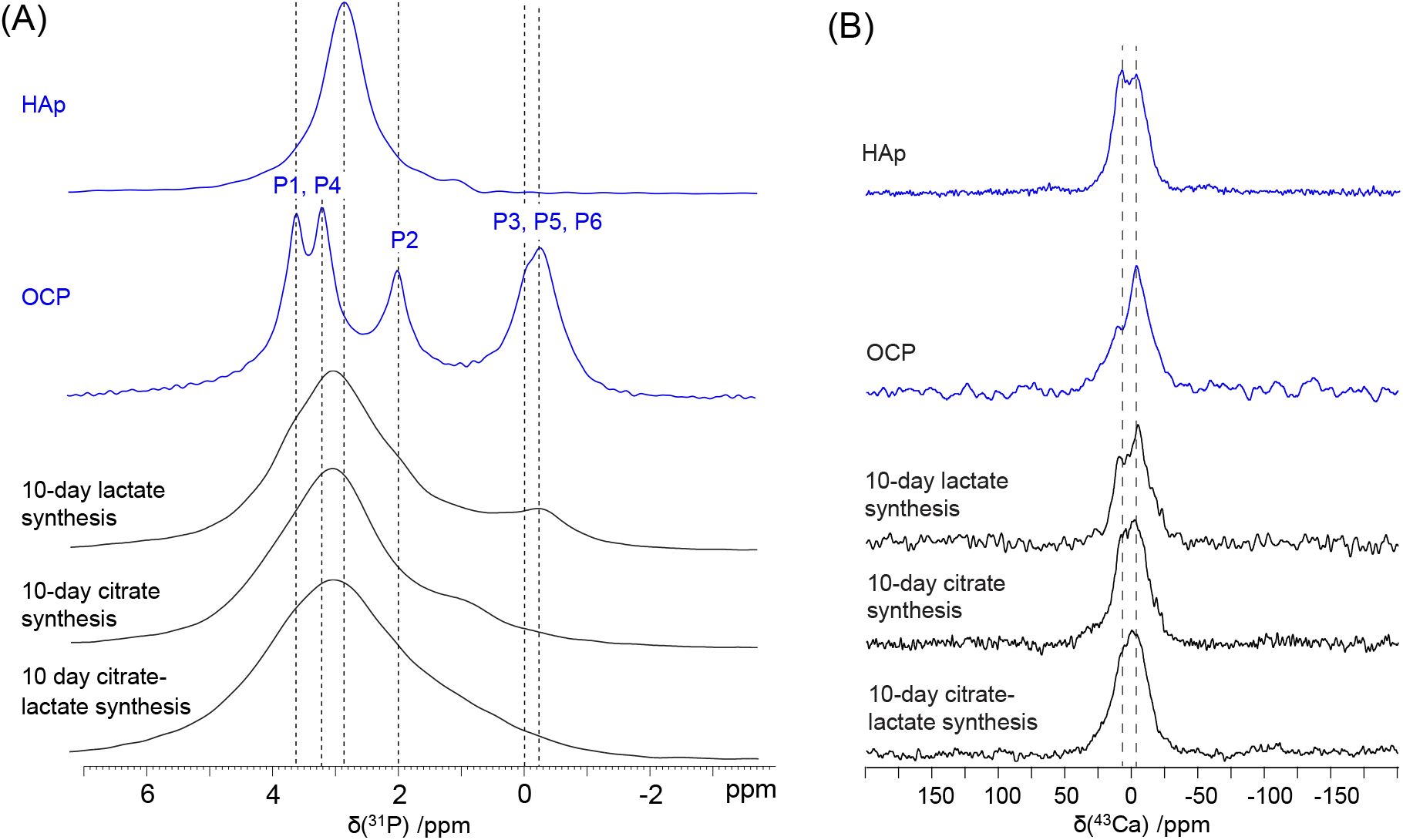
**(A)** ^31^P direct polarization (DP) MAS NMR spectra of 10-day synthesis materials investigated in this work. Sample spinning rate is 10 kHz for all samples. ^31^P DP spectra for pure HAp (nanocrystalline) and OCP are shown for reference at the top. Dotted lines indicate the chemical shifts of the characteristic HAp and OCP ^31^P signals. OCP assignments refer to the phosphate labelling in Fig 1. **(B)** ^43^Ca double-frequency sweep (DFS) NMR of the 10-day syntheses products. Spectra for pure OCP and HAp are shown for comparison. Dotted lines are for guidance only and indicate the main features of the HAp ^43^Ca spectrum.

Studies of freshwater turtles suggest that lactate too could be a component of the interfaces between apatitic mineral crystals in bone. ^10,17^ The turtle skeleton sequesters lactate and releases carbonate when the animals are in anoxic conditions, and the process reverses in normoxic conditions. This suggests that lactate can replace carbonate in bone mineral when the extracellular lactate concentration is high.^10,17^ Given the evidence for carbonate being located in disordered interfaces between apatitic mineral in bone,^15,18–20^ we hypothesized that lactate too would reside (reversibly) in bone mineral crystal interfaces.

Taking all these data together, we hypothesized that citrate, lactate and carbonate together play crucial roles in the interface structures between apatitic mineral in bone. If metabolic anions control the interfaces between mineral crystals, then it follows that they contribute to how the nanoscopic bone mineral crystals are spatially arranged and thus contribute to the 3D architecture of bone mineral. Carbonate is a necessary product of cell respiration (from carbon dioxide). Osteoblasts – bone forming cells – are professional citrate-producing cells through uptake of zinc and consequent inhibition of the tricarboxylic acid (TCA) cycle^21–23^ and the same cells can produce large quantities of lactate from glycolysis of glucose, particularly in hypoxic conditions.^24^ The presence of carbonate, citrate and lactate in bone mineral is thus inherent from the tissue metabolism and equally, aberrant levels of these metabolites can be expected as cell metabolism changes with ageing and disease. Thus understanding how these metabolites influence bone mineral structure is important for understanding the ageing process and the effects of metabolic diseases. However, as yet there is a dearth of information about how citrate and lactate can influence apatitic mineral structure. Here, we explore how citrate and lactate may contribute to bone mineral morphology and the crucial interfaces between bone mineral crystals.

## Results and discussion

Bone mineral is formed in vivo in restricted spaces bathed in extracellular fluid which contains (amongst other factors) the component mineral ions and cell metabolites. Thus, mineral nuclei form in close proximity to citrate and lactate (and carbonate) anions. To generate nanocrystalline HAp domains in a restricted space rich in citrate and lactate anions, we exploit the well-known transformation of OCP to HAp^25–31^ and the ability of OCP to incorporate carboxylic acid anions into its structure.^14,32–44^ Carboxylic acid anions locate to the hydrated layer of OCP^14,34,42,44,45^ where they bind to the apatitic-like structures either side of the OCP hydrated layers. Nucleation of HAp within an OCP lattice is hypothesized to begin in the apatitic layers of OCP and we reasoned that the presence of citrate and/ or lactate in the OCP hydrated layers would mimic the close presence of those anions around forming apatitic crystals in in vivo bone mineral formation. This mineral-only synthetic strategy allows us to decouple the effects of the metabolic acid anions from those of the collagen matrix in directing bone mineral morphology.

We synthesized OCP-carboxylic acid double salts with citrate or lactate from α-tricalcium phosphate (α-TCP; Ca_3_(PO_4_)_2_) as previously described^14,42^ with a reaction time of *48 hours* as previously optimized for OCP-citrate.^14^ The synthesis of the expected OCP-carboxylic acid double salts with approximately one metabolic acid anion per OCP unit cell (Fig 1B) was confirmed by chemical analysis (Table S1), and the OCP-double salt structure by pXRD (Fig S1), ^31^P NMR (Fig S2), ^13^C and ^13^C{^31^P} REDOR NMR (see below). We then performed the same syntheses but continued to incubate the reaction mixtures for *10 days*, to allow transformation of the initially-formed OCP-metabolic acid double salts towards apatitic mineral. The 10-day synthesis materials have similar carbon contents by wt% to the OCP-carboxylic acid double salts (see Table S1 for full chemical analyses) indicating retention of the metabolic acid anions.

We also attempted to synthesize a mixed OCP-citrate-lactate double salt, but after *48 hours* of synthesis – the same length of reaction time used to produce OCP-citrate and OCP-lactate – only small quantities of product could be detected by ^31^P NMR (Fig S2). However, maintaining the synthesis for *10 days* led to near-complete reaction of the α-TCP and a citrate-lactate-containing material with the equivalent of one citrate or two lactate anions or some linear combination thereof per OCP formula unit (Ca_16_(HPO_4_)_4_(PO_4_)_8_.10H_2_O) (Table S1).

We then used 1D ^31^P DP (Figs 1A, S2) and ^43^Ca (Fig 1B) SSNMR and pXRD (Fig S1) to confirm the formation of HAp structures in the citrate/lactate 10-day synthesis materials. The ^31^P spectra (Fig 1A) are in all cases dominated by a broad signal centred at ∼3 ppm consistent with orthophosphate in nanocrystalline HAp. Comparison of the ^31^P spectra with those for the initial double salts (Fig S2A) show similar changes to those previously observed for the transformation of pure OCP to HAp.^30^ Signal intensity in the spectral regions expected for OCP-like structures remains to some extent for all 10-day synthesis samples (> 3 ppm from OCP-like apatitic orthophosphate and around ∼0 ppm from the characteristic ^31^P signals associated with OCP-like hydrated layer PO_43-_^43-^ / HPO_4_^2-^ structures -see also Fig S2C). These are most pronounced for the lactate-only material and least for the mixed citrate-lactate material.

^43^Ca chemical shifts are highly sensitive to Ca local environments, including for bone mineral and OCP-derived phases and so are a good monitor of the mineral structures present in our materials.^33,46–48^ The ^43^Ca NMR spectrum of the lactate-only material is more similar to the one of pure OCP (Fig 1B), while those of the citrate-only and mixed citrate/lactate materials have more similarities with that of HAp (Fig 1B), in line with the conclusions made from the 1D ^31^P NMR data (Fig 1A). Interestingly, the ^43^Ca spectrum for the mixed citrate-lactate material is highly similar to that for bone, in contrast to the ^43^Ca NMR spectra nanocrystalline carbonate-substituted HAp.^56^ Specifically, the ^43^Ca spectrum for the mixed citrate-lactate material (recorded at 20 T) shows a signal maximum at –2 ppm and linewidth at half height of 1.5 kHz similar to the –3.5 ppm peak maximum and 1.4 kHz linewidth for bone recorded at 19.6 T.^47^ This suggests a similar distribution of Ca chemical environments in this mixed citrate-lactate material to those in bone, and moreover that the Ca sites in bone mineral are better modelled by the citrate-lactate apatitic material than carbonate-substituted HAp.

We next examined the HAp mineral crystal morphology in the 10-day synthesis materials with TEM. Previous studies on the transformation of pure OCP to HAp show that interlayering commonly occurs in the transformation,^30^ and thus we looked for interlayering in our 10-day materials as an indication of transformation towards HAp. TEM images for the 10-day lactate-only synthesis are dominated by nanoscopic platelets with a “picket-fence” morphology (Figs 2A, S3) with only a few particles exhibiting interlayering of electron dense and less electron-dense layers (Fig 2A, right hand side, arrow) consistent with OCP to HAp transformation. The 10-day synthesis citrate samples contain some irregular nanoscopic platelets but mainly curved nanoscopic crystals (Fig 2A and S3) exhibiting interlayering of electron dense (typically 0.9 – 1.5 nm thick) and less electron dense layers (0.9 – 1.1 nm thick) consistent with OCP to HAp transformation (Fig 2A right hand side, arrows). TEM images of the 10-day synthesis mixed citrate-lactate material show almost exclusively curved leaflet particle morphologies with interleaved fingers of electron dense HAp (1.2 – 2.4 nm thick) and less electron-dense regions (0.9 – 1.3 nm thick) (Fig 2, S4). In line with previous work^30^ and our ^31^P and ^43^Ca NMR spectra, we ascribe the electron dense interlayers to nanocrystalline HAp. The thicknesses of the less electron dense interlayers are similar for both citrate-containing materials and are consistent with the thickness of the hydrated layers in OCP and OCP-citrate. However, the ^31^P spectra in Fig 1A suggest that all molecular structures in the 10-day synthesis materials are significantly more disordered than in crystalline OCP or its double salts. Thus, we ascribe the less electron dense interlayers to disordered OCP-like hydrated layers, i.e. to regions containing disordered water and hydrogen phosphate anions, trapped between HAp structures.

**Fig 2:**
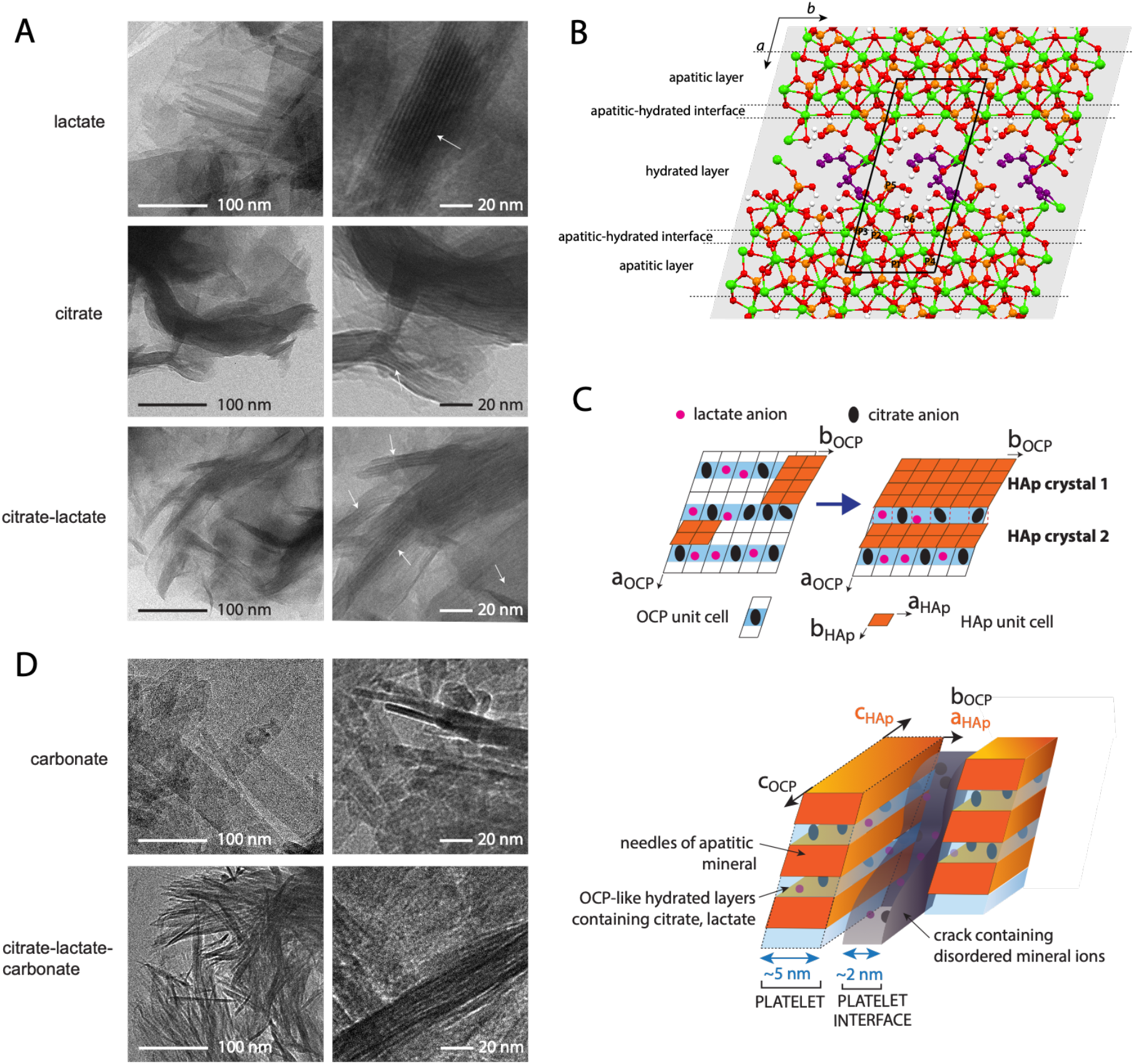
**(A)** TEM images of the 10-day synthesis samples. Arrows indicate interlayer structures. Right-hand side images are larger magnifications. **(B)** The structure of the OCP-citrate double salt determined previously by NMR crystallography^14^ showing the bridging of one citrate anion per unit cell across the hydrated layer of the OCP structure. Red, oxygen; green, calcium; orange, phosphorus; white hydrogen; purple, citrate (3-) anions. The phosphatic framework of OCP and the OCP-metabolic acid double salts (*24 hour* synthesis) consists of alternating apatitic-like (containing orthophosphate sites P1, P4) and hydrated layers (containing orthophosphate site P3 and hydrogen phosphate sites, P5, P6), with intervening apatitic-hydrated layer interface (containing orthophosphate site P2) (indicated by dotted black lines). One unit cell of the OCP-citrate structure is indicated (solid black line) and the crystallographic axes are labelled (top left). (**C) (top)** Schematic figure of the hypothesized OCP double salt – HAp transformation.^30,49^ Where transformation front meet, there is a translational mismatch in the forming HAp crystals (crystals 1 and 2 in the figure) which prevents the two crystals from coalescing. The presence of citrate/ lactate in the residual OCP-like hydrated layer stabilizes the interface between the HAp crystals. **(Bottom)** Illustration of hypothesised transformation of an initial transient OCP-citrate-lactate phase to an HAp-containing material (orange). Full transformation of the OCP phase to HAp is inhibited by the presence of citrate (black ellipses) and lactate (smaller pink circles) in the lattice, both of which need to be expelled from the lattice in order for HAp to form. Some citrate and lactate thus remain in OCP-like hydrated layers (pale blue) between regions of HAp structure. Lattice mismatch between the original OCP-like structure and the forming HAp is particularly severe along the OCP *b* axis and is expected to result in cracking perpendicular to this axis^30^ and formation of platelets (dotted line). Cracks (interplatelet gap, grey) will with ions expelled from the transforming OCP-like lattice and also likely from ions in solution forming a disordered region between HAp/ hydrated layer platelets. The expected orientation of the HAp cell axes resulting from this transformation are shown in orange, by comparison with previous work on pure OCP to HAp transformations. The platelet and interplatelet thicknesses observed in ex vivo bone are shown in blue. Note that the citrate/ lactate distributions in the figure are not intended to imply precise positions of these anions. **(D)** TEM images of samples from 10-day syntheses in which carbonate was included – see *Discussion*.

Importantly, the curved, interlayered particle morphologies found in the citrate-containing materials (Fig 1A) are similar to those found in the high-resolution studies of ex vivo bone by Reznikov et al^4^ and Xu et al.^2^ The interlayered domains in the nanoscopic platelets of our synthetic materials are resemble sideways-aggregated needles, modelling the mineral platelet morphology found in bone.^2,4^ The HAp needle/ interlayer thicknesses in our synthetic materials here are smaller than those of ex vivo bone (0.9 – 2.4 nm for the synthetic materials depending on metabolic acid composition, compared to ∼5 nm in bone^2,4^), however the size of the needles can be expected to vary with the relative and absolute concentrations of the carboxylic acid anions and with the reaction environment which is inevitably very different in bone to that in our synthetic protocol. The HAp needles in bone are curved and must be, if they are to fit the intra- and interfibrillar spaces in the bone collagen matrix.^2,4^ Here, both of our citrate-containing materials display crystal curvature. This suggests that curvature in vivo bone mineral crystals may not be entirely the result of the collagen matrix structure but may have contributions from the composition of the mineral itself.

A putative mechanism for the OCP double salt to HAp transformation, based on that deduced for pure OCP^30,49^ is depicted in Fig 2C. Importantly, the presence of metabolic acid anions in the initial OCP-double salt hydrated layers is expected to modulate the transformation from OCP to HAp. From our TEM images and from our ^31^P and ^43^Ca NMR spectra, lactate appears to inhibit transformation to HAp whilst the mixture of citrate and lactate pushes the equilibrium further towards HAp formation. This suggests that the mineral phase may be strongly influenced by the metabolic acid anion composition of the extracellular fluid in which bone mineral forms in vivo.

We next asked what are the chemical structures of the interfaces between the HAp domains in our synthetic materials and do they bear any relationship to those in bone? We hypothesized above that the interlayers between HAp domains that we observed in our materials by TEM have molecular structures resembling disordered OCP-like hydrated layers. We reasoned that since citrate and lactate are both too large to substitute into an HAp lattice, that if any citrate/ lactate remains in our 10-day synthesis materials, they will be at interfaces between HAp domains and may be involved in the OCP-like hydrated interlayers/ interfaces between HAp domains that we have deduced from ^31^P NMR (Fig 1A) and TEM (Fig 2A).

Thus, we next used ^13^C CPMAS and ^13^C{^31^P} Rotational Echo DOuble Resonance (REDOR) NMR spectra (Fig 3) to determine the chemical environment of the citrate and lactate anions in the 10-day synthesis materials to understand any interface structures that these anions are involved in. The ^13^C CPMAS spectrum for the 10-day lactate-only material (Fig 3A) is similar to that for the initial OCP-lactate double salt, the only substantive difference being modest changes in the intensity distribution for the lactate methyl signals. This similarity in ^13^C spectra between the initial OCP-lactate double salt and the 10-day synthesis material suggests relatively little change in the chemical environment of the lactate anion between the two materials, which is consistent with our conclusions from TEM and ^31^P NMR observations above that there is relatively little transformation towards HAp when lactate is present.

**Fig 3:**
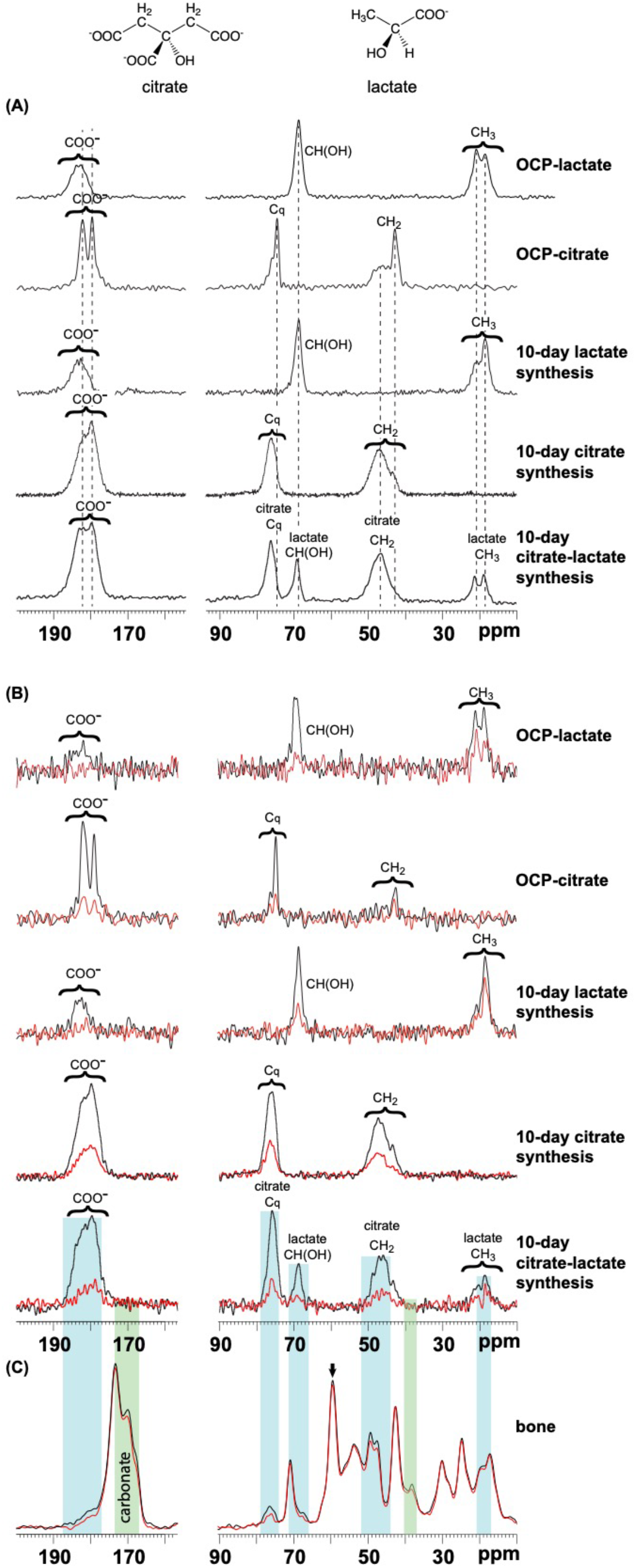
**(A)** ^13^C CP MAS spectra of the OCP-metabolic acid double salts and 10-day synthesis materials characterised in this work. Signals from citrate and lactate (structures at top) are assigned in the respective spectra. Dotted lines indicate the chemical shifts of representative ^13^C signals for the OCP-metabolic acid double salts; there are subtle chemical shift differences between these and the 10-day synthesis materials. **(B)** ^13^C{^31^P} REDOR spectra of the OCP-metabolic acid double salts and 10-day synthesis materials. The spinning rate for all samples was 10 kHz, except the OCP-citrate sample for which it was 12.5 kHz. Dephasing times are 98 1_R_ for the OCP-citrate double salt, 100 1_R_ for all other samples. Note that the low signal to noise ratio in the ^13^C{^31^P} REDOR spectrum of OCP-citrate here is due to the short ^13^C T_2_ in this material. **(C)** ^13^C{^31^P} REDOR spectrum of horse bone, dephasing time 801_R_; MAS rate 12.5 kHz. This dephasing time was chosen to minimize dephasing on collagen signals in order to highlight the dephasing on non-collagen signals. Coloured bars indicate the main spectral regions with REDOR dephasing; blue bars are regions where there is also REDOR dephasing in the spectrum for the mixed citrate-lactate synthetic material in (B); green bars are regions where there is no corresponding dephasing in the mixed citrate-lactate material spectrum. The expected region for carbonate ^13^C signals (and thus REDOR dephasing due to the presence of carbonate in bone mineral) is indicsated. Note that the bone spectrum here is dominated by signals from collagen and that any signals from metabolites in the bone mineral are expected to be very small in comparison. Thus, REDOR dephasing may occur in spectral regions between the dominant signals, such as in the 44 – 47 ppm region. Equally, signals from collagen are expected to also dephase to some extent as bone mineral is in proximity to collagen.

There are small changes in the ^13^C chemical shift distribution for the citrate-only and mixed citrate-lactate 10-day synthesis materials compared to the OCP-citrate and OCP-lactate double salts (Fig 3A) indicating only subtle changes in the chemical environment of the citrate/lactate in the 10-day citrate-containing materials compared with those in the OCP double salts. The citrate and lactate anions are contained in hydrated layers between apatitic layers in the OCP double salts. The disordered OCP-like hydrated interlayers between the HAp domains observed by TEM and ^31^P NMR for these materials represents a similar chemical environment and hence we propose that the ^13^C signals for the 10-day synthesis HAp-containing materials come primarily from citrate/lactate residing in these disordered, hydrated interlayers/ interfaces between HAp domains. We note that some small residual signal from untransformed OCP-citrate is expected in the case of the citrate-only material, for which ^31^P and ^43^Ca NMR and TEM shows the transformation to apatitic material is not fully complete.

This hypothesis is confirmed by ^13^C{^31^P} REDOR spectra (Fig 3B). In the REDOR spectra, spatial proximity of the metabolic acid anions to phosphate is identified by reduction of anion’s ^13^C signal intensity between the reference and REDOR spectra. Despite the inevitable low signal-to-noise ratio of the REDOR spectra (because of low net carbon content and *T*_2_ losses), it is clear that all ^13^C signals for the 10-day synthesis materials exhibit a substantial REDOR dephasing effect which is at least of similar order of magnitude to those for the OCP double salts (Fig 3B). The magnitude of the REDOR effect is greater the more phosphate sites there are around the metabolic acid anion and the closer together the anion and phosphate are. Citrate and lactate reside unequivocally in the hydrated layers within the OCP lattice structure of the 48-hour synthesis OCP double salts and so the REDOR effect for these double salts comes from citrate and lactate fully surrounded by close-proximity phosphate ^31^P. The magnitude of the REDOR effects for the double salts can therefore be thought of as an upper limit on the extent of REDOR effect that can possibly occur. That the REDOR effect on all the metabolic acid ^13^C signals for the 10-day synthesis materials is at least as great as for the signals for the corresponding double salts suggests that the metabolic anions in the 10-day synthesis materials must also be fully surrounded by phosphate. In the case of the lactate-only material, that is clearly because there has been relatively little transformation away from the initial OCP-lactate double salt, but the citrate-only and mixed citrate-lactate materials exhibit considerable extents of transformation towards HAp, particularly for the latter material. Thus, the REDOR effects for the 10-day citrate-containing materials come primarily from metabolic acid anions in proximity to HAp layers, not the original OCP structure. If the metabolic acid anions resided on single 2D HAp surfaces (as opposed to between two HAp surfaces), the number of nearby phosphate ions would inevitably be lower and so the REDOR effect would be smaller. If one takes the model that the metabolic acids reside between two apatitic layers in the OCP double salts, then binding to a single apatitic layer can be expected to yield approximately half the size of REDOR effect, and this is clearly not the case. Thus, we conclude that the metabolic acid anions in the 10-day synthesis citrate-only and mixed citrate-lactate materials are between apatitic-like layers/ surfaces, consistent with the metabolic acid anions in these citrate-containing materials being in the interlayers between the HAp layers.^†^Bone mineral contains both citrate and lactate so we then explored how the citrate and lactate environments the mixed citrate-lactate 10-day apatitic material compare with those in bone mineral by comparing the REDOR spectra for the respective samples. The ^13^C NMR spectrum of bone is dominated by signals from collagen; bone mineral is intimately associated with collagen in bone and so collagen ^13^C signals are expected to dephase at least to some extent in ^13^C{^31^P} REDOR experiments. The high concentration of collagen in bone means that REDOR dephasing of collagen ^13^C signals can easily obscure any REDOR effect from the metabolites in bone mineral. Thus, we chose a REDOR dephasing time for the bone sample where the REDOR dephasing on collagen signals is minimal to emphasize the spectral regions where there is dephasing of ^13^C signals from metabolic acid anions. The regions of the bone spectrum in Fig 3C that suffer the most extensive REDOR dephasing (reduction in intensity between the reference and REDOR ^13^C spectra) are highlighted with coloured bands. Blue bands are spectral regions where there is also REDOR dephasing in the corresponding region of the spectrum for the mixed citrate-lactate apatitic material; green bands are where there is no such corresponding dephasing. As can be seen from Fig 3C, a substantial part of the bone spectrum dephasing may be accounted for by citrate and lactate environments similar to those in the synthetic citrate-lactate material and a further part by the known incorporation of carbonate into bone mineral.

Thus, the disordered OCP-like hydrated interfaces between the HAp domains in our synthetic apatitic materials appear to be good models of the chemical environment of citrate and lactate in bone mineral. The interfaces between the sideways-aggregated apatitic needles of bone mineral are the obvious analogues of the citrate/lactate-containing interlayers between the HAp needle-shaped domains/ interlayers of our synthetic materials. Thus, we propose that the sideways aggregated needles in bone mineral are supported by citrate/ lactate-containing interfaces.

In addition to the interfaces between the sideways-aggregated apatitic needles in bone mineral, there must also be chemical interfaces between the resulting aggregate platelets as the platelets typically remain bound together even if the underlying collagen matrix is removed [refs]. The TEM images of our synthetic materials showed that the interlayered platelets in them are typically aggregated into multiple layers of platelets which we could not disperse without fracturing the platelets, implying that there are relatively strong interface structures aggregating the platelets together. Thus, we next used 2D ^1^H-^31^P correlation NMR spectroscopy to garner information on the molecular structures involved in these inter-platelet interfaces.

2D ^1^H-^31^P correlation spectra allow the molecular structures of biominerals to be characterized by defining the spatial proximity of phosphate/ hydrogen phosphate anions and ^1^H-containing components of the mineral, namely water and hydroxyl groups (and hydrogen phosphate anions). Signals are readily observed in the 2D spectra from disordered structural regions as well as more ordered regions, making these spectra highly useful for assessing the molecular structures in interfaces between HAp domains.^15,16,46,50–56^ A 2D ^1^H-^31^P correlation spectra of the 10-day synthesis citrate-lactate material is shown in Fig 4A and it exhibits the expected correlation signal for HAp at ^31^P 3 ppm – ^1^H 0 ppm. Importantly, this is the only expected signal from HAp; all the remaining signals in the 2D spectrum are from non-apatitic components, namely the interfaces between HAp domains, and so we anticipate signals from both the interlayers between HAp needle-shaped domains within mineral platelets and the interfaces between the mineral platelets. We can readily assign the ^31^P-^1^H correlation signals from the citrate/lactate interlayers between HAp needles as they can be expected to have ^1^H-^31^P chemical shifts similar to those in the OCP double salts. The 2D ^1^H-^31^P spectrum for the lactate-only 10-day synthesis material in Fig SX exemplifies the expected ^1^H chemical shifts for the citrate/lactate-containing interlayers as the lactate-only material we know is predominantly still the OCP-lactate double salt structure. The ^1^H shifts for the OCP-citrate double salt have also been previously published [ref]. Thus, we know that ^1^H chemical shifts around ∼6 ppm are from water in the interlayers/ interfaces between HAp needles and ^1^H chemical shifts around ∼14 ppm are from HPOanions in those same interfaces.

**Fig 4:**
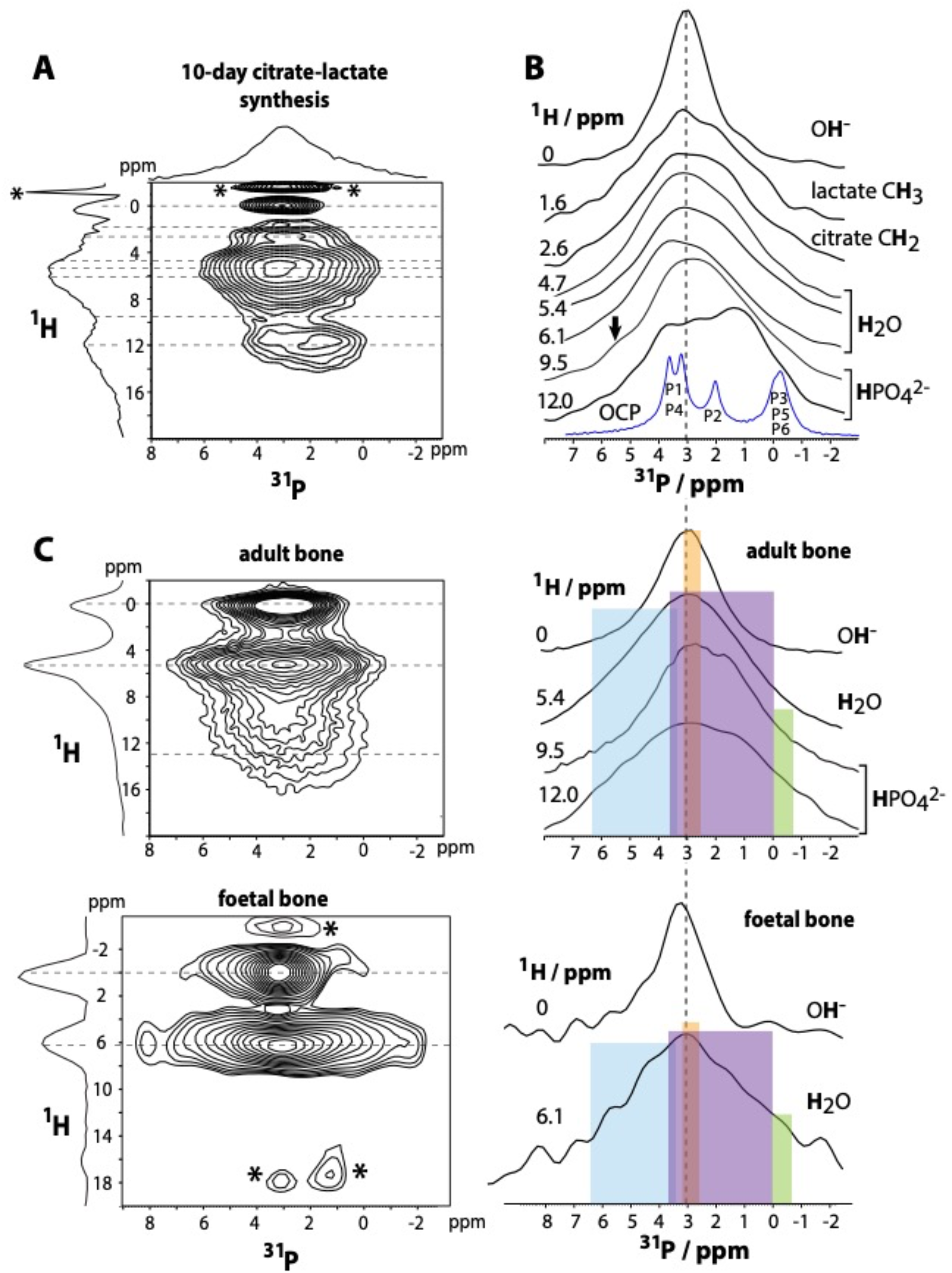
**(A)** 2D ^1^H-^31^P heteronuclear correlation spectra of the 10-day citrate-lactate synthesis. * indicate spectral artefacts from sinc wiggles. Dotted lines indicate the slices from which ^31^P spectra are taken in (B). **(B)** ^31^P slices taken from the 2D ^1^H-^31^P heteronuclear correlation spectra in (A). The ^1^H chemical shifts at which the slices are taken are indicated on the left, and the assignment of those ^1^H chemical shifts are shown on the right. Arrows mark spectral features referred to in the text. **(C)** 2D ^1^H-^31^P heteronuclear correlation spectra for adult sheep bone and foetal bone growth plate (sheep). Note that the foetal bone sample is necessarily very small and dehydrates rapidly in the spectrometer leading to low signal to noise ratio; the signals marked with asterisks here are noise. **(D)** ^31^P slices taken from the 2D ^1^H-^31^P heteronuclear correlation spectra in (C). The ^1^H chemical shifts at which the slices are taken are indicated on the left, and the assignment of those ^1^H chemical shifts are shown on the right. The absence of a HPO_4_^2-^ signal for the foetal bone growth plate may be due to low signal to noise ratio for this spectrum. The chemical shift ranges expected for different ^31^P species are indicated for the bone samples by: orange, HAp PO_4_^3-^; purple, OCP PO_44_^3-^; blue, HAp surface HPO_4_^2-^; green, OCP HPO_4_^2-^ (see Fig S3 for further details). The ^31^P (direct polarization) spectrum is shown for comparison of the ^31^P slices throughout. The vertical dotted line allows comparison of the ^31^P chemical shift for adult bone mineral orthophosphate with the ^31^P spectrum of the synthetic material and the foetal bone spectrum.

We then reasoned that any ^1^H signals and associated correlated ^31^P signals outside these ranges may be assigned to water and HPO_4_^2-^ in the interplatelet interfaces as the only other significant phosphatic chemical environment likely to be present in our synthetic citrate-lactate-containing apatitic material.

The 2D ^1^H - ^31^P spectrum exhibits significant signal intensity at lower water ^1^H chemical shifts e.g. 4.7 ppm in Figs 4A,B, in addition to the OCP-like interface water ^1^H signal ∼6 ppm. Water ^1^H chemical shifts in the range ∼4.5 – 5 ppm are consistent with water on HAp nanocrystal surfaces^15^ and indeed the 4.7 ppm ^1^H signal in Figs 4A,B correlates with a broad HAp orthophosphate ^31^P signal at ∼2.9 ppm as would be expected for water on an HAp surface. The lactate methyl ^1^H signal at (Fig 4A) correlates with similar ^31^P spectrum to the HAp surface water 4.7 ppm ^1^H signal, suggesting that lactate and HAp external surface water “see” similar phosphatic environments, and thus that the interplatelet interface contains some lactate.

The HPO_4^2-1^_H signal contains chemical shift components (9 – 11 ppm, Figs 4A, B) that too low to be from OCP-like hydrated interlayers but which are consistent with HPO_4^2-1^_on HAp surfaces^15,55–57^ and indeed correlate with a broad HAp ^31^P signal at ∼ 3 ppm (Figs 4A, B), and with a putative broad shoulder at ∼5.5 ppm (Fig 4B, 9.5 ppm ^1^H slice, arrow) which is within the expected ^31^P chemical shift range for HPO_4^2-1^_on HAp surfaces.^55,56^ Thus, the HAp external surfaces in our synthetic material have a disordered hydrated structure common to nanocrystalline HAp. Importantly, we can then conclude that the interlayered mineral platelets of the citrate-lactate synthetic material have *interplatelet interfaces* containing disordered, hydrated HPO_4^2-1^_with both water and HPO_4^2-1^_anions in distinctive chemical environments that are different to those between the HAp domains within the mineral platelets, which are organized by the citrate/ lactate anions.

We then compared the HAp interface structures as observed by 2D ^1^H-^31^P correlation NMR spectroscopy for the 10-day synthesis mixed citrate-lactate material with those for adult (mature) and developing (foetal growth plate) bone. The 2D ^1^H-^31^P heteronuclear correlation spectrum of mature bone (Fig 4C) shows the expected signal from nanocrystalline HAp structures, namely the (broad) correlation signal at ^31^P ∼3 ppm – ^1^H 0 ppm. Additional signals from highly hydrated, disordered ortho-phosphate and hydrogen phosphate anions have been observed previously, and assigned to disordered non-apatitic material on the surfaces of HAp domains.^15,16,20,46,50–54^ The spectrum for the 10-day mixed citrate-lactate material (Figs 4A, B) shows strong similarities to that for mature bone (Figs 4 C, D, top)): the ^31^P ∼3 ppm – ^1^H 0 ppm apatitic signal, the water ^1^H signal with chemical shift maximum at 5.5 ppm compared with 5.4 ppm in adult bone (in the context of a range of ∼4.8 - ∼6 ppm for water in calcium phosphate minerals^56^) and the distribution of hydrogen phosphate ^1^H chemical shifts.^20^ The ^31^P spectra correlating with the 5.4 ppm water ^1^H and 9.5 ppm HPO_4^2-1^_H signals for bone (Fig 4D, top) show strong similarities in chemical shift range and intensity distribution to those for the mixed citrate-lactate material (Fig 4B). The ^31^P spectrum correlating with the 12 ppm HPO_42-1_H signal for the mixed citrate-lactate material (Fig 4B) again covers a similar chemical shift range as for the equivalent signal for adult bone, although there are resolvable signals in the ^31^P spectrum for the synthetic material (Fig 4A) but not for the bone sample (Fig 4C), indicating a more heterogeneous distribution of structures for HPO_42-_ sites with this ^1^H chemical shift in native bone compared to the synthetic material. This is to be expected when one considers that a native bone sample contains mineral deposited over a continuum of time points and shaped at least to some extent by the underlying collagen matrix, neither of which points are modelled in our synthetic material. Our synthetic mixed citrate-lactate material contains significantly more interface structures between apatitic domains than bone mineral because the apatitic domains in our synthetic material are smaller than in bone mineral. Thus, we would expect the signals for the interface regions, i.e. ^1^H-^31^P correlations involving water and HPO_4^2-^_, to be relatively more intense for our synthetic material. Given these constraints, there is remarkable homology between the 2D ^1^H-^31^P spectra for our mixed citrate-lactate material and adult bone mineral.

Intriguingly, the 2D ^1^H-^31^P NMR spectrum of foetal bone growth plate (Figs 4C, D, bottom) shows more similarity to those from OCP-like phases: the OH^-1^H – PO_4 ^3-31^_P apatitic signal (Fig 4C, bottom) is centred at a ^31^P chemical shift of 3.25 ppm, which is more indicative of OCP-like apatitic orthophosphate environments than nanocrystalline hydroxyapatite orthophosphate. The OH^-1^H – PO_4 ^3-31^_P correlation signal is also much broader in the ^31^P spectral dimension for the foetal bone growth plate sample than the corresponding signal from the adult bone sample, indicating that the apatitic orthophosphate component in foetal bone is more structurally disordered than in adult bone. The water ^1^H signal (Fig 4C, bottom) is centred at 6.1 ppm for foetal bone rather than 5.4 ppm in adult bone. This water ^1^H chemical shift is more similar to that for water in OCP-like structures^14,58,59^ than for nanocrystalline HAp surface water (4.85 ppm).^55,56^ The water ^1^H signal in the foetal bone spectrum is correlated with a wide range of ^31^P chemical shifts that includes the ranges expected for HAp and OCP-like orthophosphate, Hap surface HPO_4 ^2-^_ and OCP-like HPO_4 ^2-^_ (Fig 4D, bottom).^55,56^ There is no clear HPO_4 ^2-1^_H signal from the foetal bone growth plate however (Fig 4C, bottom). This may be because of the inevitable low signal-to-noise ratio for this spectrum – there is an inherently small sample volume (the bone growth plate region is very small) and there is limited possible experiment time because of degradation of the foetal bone sample in the spectrometer. The ^1^H-^31^P NMR spectrum of foetal bone here only gives a single snapshot of the mineral phase transformations that may occur in vivo, but the presence of both disordered apatitic orthophosphate groups which are more OCP-like than HAp-like in terms of ^31^P chemical shift, and which are associated with OCP-like water environments in this sample suggest a process of bone mineral development occurring through one or more OCP-like intermediate phases. This is not the first time that an OCP-like phase has been suggested as an intermediate phase in the formation of bone mineral^60,61^ but the possibility is still contentious.^18,19^

Whether the HAp domains in bone form through an intermediate OCP-like lattice or directly within an amorphous calcium phosphate (ACP) phase that is widely believed to be the first-formed mineral in bone,^2,60,62–64^ the HAp nuclei in bone form within a restricted space and within a highly complex chemical environment. That chemical environment necessarily includes osteoblast and blood plasma metabolites including carbonate, citrate and lactate, and the key questions here are: where do these metabolites reside in the mineral structure and what do they contribute to that structure? Carbonate is known to substitute readily for both orthophosphate and hydroxyl anions in HAp, including the apatitic regions of bone mineral.^11,15,20,65,66^ In synthetic HAp materials at least, carbonate plays a role in controlling the size of HAp domains through the lattice defects its incorporation causes, typically resulting in nanoscopic HAp crystals.^15,56,67^

HAp forming in the presence of carbonate is likely to contain carbonate substitutions, whatever the route of its formation. Carbonate-substituted OCP^68,69^ has already been shown to hydrolyze to carbonate-substituted HAp^69^, but we briefly explored the mineral morphology effects of carbonate substitution in our current synthetic platform. Figure 2D shows TEM images of the morphology of mineral generated from the transformation of carbonate-containing OCP using an identical synthesis as for the citrate/lactate-containing materials. ^31^P NMR spectra (Fig S2) show the material to be a mixture of HAp and OCP structures, similar to the lactate-only 10 day synthesis material. The TEM images (Fig 2D) show a mixture of mainly tabular crystals and a smaller number of partially merged more needle-like crystals.

Then, as a final step in this work, we briefly examined whether the combination of carbonate-citrate-lactate may have any effect on mineral morphology above that we have already found for citrate-lactate. Figure 2D shows TEM images from syntheses of apatitic mineral using the same approach of transformation from metabolic acid anion-containing OCP from OCP synthesized with carbonate, citrate and lactate. The mineral in this case consists of aggregated needles, most with some curvature, not dissimilar to the citrate-lactate-containing material, but the needle diameters are typically larger than for the citrate-lactate material.

In contrast to carbonate, citrate and lactate anions are too large to substitute for HAp anions. However, as the transformation of OCP-carboxylic acid double salts to HAp characterized in this work demonstrates, forming HAp crystals within a restricted space (here within the lattice of nanocrystalline OCP) in the presence of citrate and lactate leads to the inclusion of both anions in the resulting material. Citrate and lactate in our synthetic system formed hydrated interfaces between HAp domains and inhibited fusion of growing HAp crystals into larger crystals. It has previously been proposed that monolayers of calcium citrate tetrahydrate can form preferentially between some HAp crystal faces.^70^ The authors of that study^70^ proposed an intriguing model in which such monolayers forming between preferential HAp crystal faces cause specific aggregation patterns of HAp crystals, and suggested that such a model may account for the 3D architecture of bone mineral. The sideways aggregated HAp needle morphology of bone mineral insists that something “glues” neighbouring HAp needles together through the appropriate HAp surfaces. The distance between the HAp needles within a bone mineral platelet was estimated to be less than 1 nm,^4^ which strongly suggests that the “glue” must consist of relatively small molecules and not HAp-binding proteins, for instance. The citrate/ lactate-containing hydrated interlayers in our 10-day synthesis apatitic citrate-only and mixed citrate-lactate materials fit the less than 1 nm thickness condition for the inter-HAp needle region. Our experimental data is consistent with the HAp needles in the mixed citrate-lactate material having two different types of surface: “internal” surfaces with hydrated, citrate-containing interlayers between HAp needles and external surfaces containing water and hydrogen phosphate similar to those on synthetic nanocrystalline HAp.^15,18,19,46,50,52,53,71^ Where bone mineral platelets stack together, these latter external surfaces will form the interplatelet interfaces. It seems likely that metabolic anions are also present in these interplatelet interfaces. The 2D ^1^H-^31^P NMR data for our mixed citrate-lactate material suggest the lactate anion in this material is substantially present in the interplatelet regions. Carbonate has previously been shown to be present in the non-apatitic structures in bone mineral,^15,20^ though whether these are the inter-HAp needle regions within the mineral platelets or the regions between the resulting platelets or both is not yet clear.

Citrate appears to be important for crystal curvature; both the citrate-only and mixed citrate-lactate materials exhibit crystal curvature, whilst the lactate-only and carbonate-only materials (Figs 2 and S2) and carbonate-containing nanocrystalline HAp do not.^15^ We speculate that the cause of crystal curvature is that citrate can adopt many different conformations in the interface between HAp nanocrystals/ domains and thus span different distances between apatitic surfaces.^14^ Regions of varying interlayer thickness between two HAp layers will cause curvature of the layers if the size dimensions of such regions are significant compared to the overall layer dimensions and a similar argument can be extended to stacks of more than two HAp layers.

Lactate inhibited the transformation towards HAp in our model materials here. Hence, we hypothesize that high lactate concentrations may inhibit formation of HAp in vivo as well. Intriguingly, foetal bone contains very high quantities of lactate compared to citrate perhaps as a result of the more hypoxic environment within foetal tissues than in mature tissues.^13^ The 10-day synthesis lactate-only material is a soft, chalky like substance, very different to the materials containing citrate which are much harder, properties which we confirmed with AFM (Fig S8). Foetal bones are necessarily soft to allow for the birth process and we propose that the high lactate content of foetal bones is necessary to keep the bones soft until postpartum and that lactate achieves this by inhibiting the transformation towards HAp. If citrate is important in governing bone mineral 3D architecture, we would expect that citrate is also important for bone mechanical properties and early evidence for this comes from osteoporotic mice which were found to have significantly lower bone citrate content than in controls.^75^

Increasing prevalence of bone pathologies associated with ageing and metabolic diseases have highlighted connections between bone cell metabolism and the molecular structure of bone mineral. A model in which the 3D architecture of HAp nanoregions in bone is governed by metabolic-acid-rich material between specific surfaces of HAp intergrowths can potentially explain how bone mineral 3D architecture and mechanical properties are influenced by bone cell and whole animal metabolism. The production of both citrate and lactate in bone is in part determined by the glucose and oxygen supply^7,8,72^ and partly by a delicate balance of hormones and growth factors known to be important in bone health^7,24,73^ including insulin, estradiol^7^ and parathyroid hormone. The phase diagram for calcium phosphate-carbonate-citrate-lactate-water is undeniably complicated and it is possible that even subtle changes in relative and absolute concentrations of carbonate, citrate and lactate in vivo result in significantly different size distribution of HAp regions and composition and spatial arrangement of non-apatitic regions – with significant consequent implications for the bone mineral mechanical properties. Reduction of citrate/ lactate concentrations during bone mineral formation would be expected to lead to larger HAp domains, based on our findings here. Bone HAp crystals are known to increase in size and crystallinity (atomic order) with animal ageing, regardless of the amount of mineral or collagen per unit volume of the bone tissue, and the increasing size and crystallinity correlates with increasing bone fragility.^74^ This important phenomenon has yet to be explained; importantly, there is no established correlation of mineral fragility with Ca/P ratio or carbonate content. Indeed, the carbonate environment appears not to change with age.^66^ We hypothesize that this bone-ageing characteristic results from complex changes in carbonate/citrate/ lactate concentrations when bone is remodelled, due to inevitable changes in osteoblast and/ or whole-body metabolism with ageing.

There are many other calcium-binding metabolites that could also be present during bone mineral formation, and which may also affect bone mineral structure. The bone REDOR spectrum in Fig 5C shows two spectral regions of dephasing that cannot be accounted for by citrate or lactate. One region is in the chemical shift range for carbonate (Fig 5C), but the other (38 – 40 ppm) suggest that another metabolite may also be present in bone mineral. We have previously shown that OCP citrate^14^ has a ^13^**C**H_2_ signal in this spectral region (see also Fig 5A) and it is possible that alternative citrate environments in bone mineral may account for this REDOR dephasing. An interesting possibility is the incorporation of pathological metabolites into bone mineral, pathological in the sense that their incorporation drives aberrant mineral architectures. We have previously shown that succinate, another product of the TCA cycle, under the same reaction conditions as those used in this work, forms an OCP-succinate double salt which is highly crystalline, forming large crystals compared to OCP-lactate and OCP-citrate, with ordered succinate anions bridging the OCP hydrated layer.^42^ Whilst elevated temperatures induce transformation of OCP-succinate to HAp,^49^ incubation in our reaction conditions at 37ºC for 10 days did not. This suggests the intriguing possibility that some metabolic anions may be pathological to bone mineral formation in that they inhibit the necessary phase transformations and stabilize pathological mineral structures. In turn, this suggests that the dysregulation of osteoblast and whole-body metabolism prevalent in ageing, could be a significant factor in pathologies associated with bone ageing.

## Conclusions

From our work here, we propose that the intricate 3D architecture of bone mineral^4^ requires the presence of multiple metabolic acid ions during mineral formation. The presence of the metabolic acid anions, citrate, lactate and carbonate, results in the formation of complex nanocrystalline apatitic phases where HAp nanocrystals interface through disordered, hydrated metabolic anion-containing interfaces. The presence of citrate generates curved mineral crystals and this may facilitate the shaping of bone mineral into the niches of the underlying collagen matrix. We have demonstrated here how citrate and lactate affect transformation to HAp, lactate favouring the initial OCP phase of our experiments, and a mixture of citrate and lactate favouring transformation to particles with interlayered HAp/ disordered citrate/ lactate-containing regions that model the sideways-aggregated HAp needles of in vivo bone mineral. We hypothesise that other cell metabolites may also be involved in forming the 3D architecture of bone mineral in vivo and thus the mechanical properties of bone mineral. A model in which cell metabolites control the 3D bone mineral architecture and calcium phosphate composition can potentially explain why bone mineral properties change in ageing and pathologies such as osteoporosis. Further investigations on the effects of other cell metabolites on bone mineral structure are thus urgently needed.

## Acknowledgements

YL was funded by an EPSRC doctoral training program studentship. RL was funded by a Cambridge Trust and China Scholarship Council. The UK High-Field Solid-State NMR Facility used in this research was funded by EPSRC and BBSRC (<u>EP/T015063/1</u>), as well as the University of Warwick including via part funding through Birmingham Science City Advanced Materials Projects 1 and 2 supported by Advantage West Midlands (AWM) and the European Regional Development Fund (ERDF). TEM and SEM images were recorded at the Cambridge Advanced Imaging Centre (CAIC).

## Materials and Methods

### Synthesis

#### OCP-citrate double salt

A solution of citric acid (2.643 g, 13.8 mmol) in HPLC-grade water (55 mL) was maintained at 37 ºC, and the pH adjusted to 5.52 by dropwise addition of conc. NaOH solution. α-TCP (836.1 mg, 2.70 mmol) was then added, and the pH of the resulting suspension adjusted to 6.50 by dropwise addition of conc. NaOH. The suspension was stirred at 200 rpm for 48 hours, after which the solid was isolated by gravity filtration and dried in air overnight at room temperature. The final product was a white powder (215 mg, 0.1 mmol)

#### OCP-lactate double salt

Sodium-L-lactate (3.64g, Sigma-Aldrich) was dissolved in 130 ml distilled water maintained at 37 ºC and then α-TCP (1.04g, Sigma-Aldrich) was added while stirring. The pH was adjusted to 6.50 by dropwise addition of concentrated HCl solution. The reaction mixture was covered and stirred at 37°C, 500 rpm for 24 hours after which the product was isolated by gravity filtration, washed with distilled water and dried in air overnight at room temperature.

The hydrolysed OCP citrate and OCP-lactate were synthesised as above except that the reaction mixture in each case was stirred at 300 rpm for 10 days (at 37°C)

#### OCP-citrate-lactate

A solution of citric acid (1.32 g, 6.9 mmol) and sodium L-lactate (1.4g, 12.5 mmol) in distilled water (100 ml) was maintained at 37 °C, and the pH was adjusted to 5.50 by dropwise addition of NaOH. α-TCP (1.60 g, 5.3 mmol) was then added and the pH was adjusted to 6.50 by dropwise addition of NaOH. The suspension was stirred at 300 rpm for 10 days, and the solid was collected by gravity filtration and dried in air.

C,H elemental analysis was perfomed with an Exeter Analytical CE440 elemental analyser with combustion at 975°C. Ca and P elemental analyses were measured with a Thermo Scientific 7400 ICP-OES instrument at 396.84 nm and 178.28 nm respectively.

### Solid-state NMR (SSNMR) spectroscopy

A Bruker 400 MHz Avance spectroscopy II spectrometer was used for solid-state ^1^H, ^13^C and ^31^P NMR measurements, at frequencies of 400.42 MHz, 100.6 MHz and 162.1 MHz respectively, with standard Bruker double and triple resonance, MAS probes. Samples were packed into disposable HR-MAS inserts where necessary, and loaded into the 4 mm zirconia rotors for magic angle spinning (MAS), at a rate of 10 kHz, unless otherwise stated.

Samples were characterised using ^31^P direct-polarisation (DP), typically using a 2.5 μs 31P (90°) and 600 s recycle delay. ^31^P {^1^H} CP spectra for the 48-hour mixed citrate-lactate samples were recorded with ^1^H 90° pulse length, 2.5 μs, ^31^P 90° pulse length, 2.57 μs, ^1^H-^31^P CP contact time, 10 ms and recycle time, 2 s. REDOR experiments used typical REDOR dephasing times of 2-10 ms, corresponding to 20 – 100 ι−_R_ (100 ι−_R_ for spectra in Fig 4, except as in figure caption). Broadband TPPM decoupling during signal acquisition was used in all ^13^C and ^31^P experiments.

_1_H-^31^P heteronuclear correlation experiments were performed with Frequency-Switched Lee-Goldburg (FSLG) decoupling during t_1_ (^1^H field strength 100 kHz, 2 ms contact time, 1 s recycle delay). Broadband SPINAL 64 decoupling was used during t_2_ signal acquisition.

_13_C spectra were referenced to the glycine C_α_ signal at 43.1 ppm relative to TMS at 0 ppm. ^31^P spectra were referenced to the hydroxyapatite ^31^P signal at 2.85 ppm relative to 85 wt% H_3_PO_4_ at 0 ppm. ^1^H spectra were referenced to the hydroxyapatite signal at 0 ppm relative to TMS at 0 ppm.

OCP and OCP-metabolic acid materials were studied using natural abundance _43_Ca NMR experiments, which were acquired at 20.0 T on a Bruker Avance III-850 (850 MHz ^1^H frequency) spectrometer at the UK 850 MHz Solid-State NMR Facility, with the assistance of Dr Dinu Iuga, operating at ^43^Ca Larmor frequency of 57.22 MHz, using a low-γ 7 mm Bruker MAS probe spinning at 5 kHz. For the simple OCP phase, a RAPT (rotor assisted population transfer) enhancement scheme was used (offset of 150 kHz, RF ∼ 9 kHz), followed by a 90° selective solid pulse of 1.5 μs. A total of 137864 transients were acquired, with a recycle delay of 0.5s. For all the intercalated OCP samples, a multi-DFS (double frequency sweep) enhancement scheme followed by a 90° selective pulse of 1.5 μs, was used,^1,2^ which was first optimized on a ^43^Ca-labeled CaHPO_4_ sample.^3^ A total of 65500 transients were acquired, with a recycle delay of 0.5s. A ^43^Ca-enriched HAp phase was also studied at 20.0 T for comparison to OCP phases. In this case, a Bruker 4 mm probe was used, and 64 transients were acquired, with a recycle delay of 0.8 s. All ^43^Ca chemical shifts were referenced at 0 ppm to a 1 mol.L^‒1^ aqueous solution of CaCl_2_.^4^

### Powder X-ray diffraction

PXRD experiments were performed on a Philips X’Pert Pro powder diffractometer equipped with an X’celerator RTMS detector and using Ni-filtered Cu Kα radiation of wavelength 0.154 nm. Samples were ground into fine powders and mounted on flat steel plates and data collection was performed over the range 2θ = 3 to 60°. Different amounts of sample were used for the different compounds and hence the signal-to-noise ratio of the diffraction patterns differ between samples.

### TEM

TEM images were taken using a Tecnai G2 80 – 200 kV transmission electron microscope (Cambridge Advanced Imaging Centre) operating at 120 or 200 kV.

## Footnotes

† We note that the relative intensities of the ^13^C CPMAS NMR signals from lactate in the 10-day synthesis mixed citrate-lactate material are lower than those of the citrate ^13^C signals for this material (Fig 3A). The lower relative intensity of the lactate ^13^C signals may reflect lower relative incorporation of lactate anions compared to citrate and/ or that these smaller anions are dynamic on the NMR timescale in this structure, resulting in lower cross-polarization intensity. The lactate C-OH and CH_3_signals in the ^13^C{^31^P} REDOR reference spectrum of the mixed citrate-lactate material (Fig 3B) have even lower relative intensity compared to the citrate signals than in the ^13^C CPMAS spectrum for this material. Thus, the ^13^C T_2_ for the lactate ^13^C signals in this material must be shorter than for the citrate signals. Shorter ^13^C T_2_ is consistent with the lactate anions in this material having a greater degree of millisecond timescale molecular motion than citrate anions in this material, as suggested above when assessing the lower CP intensity for the lactate component in this material. Thus, we conclude that the lower lactate ^13^C signal intensity compared to citrate is at least in part due to molecular dynamics of the lactate which restricts the ^13^C CP intensity generated and does not necessarily indicate a lower proportion of lactate in this mixed citrate-lactate material.

1 K. M. N. Burgess, F. A. Perras, I. L. Moudrakovski, Y. Xu, D. L. Bryce, *Can. J. Chem.* 2015, 93.

2 A. Brinkmann, A.P.M. Kentgens, *J. Phys. Chem.* B, 2006, 110, 16089.

3 C. Bonhomme, X. Wang, I. Hung, Z. Gan, C. Gervais, C. Sassoye, J. Rimsza, J. Du, M. E. Smith, J. V. Hanna, S. Sarda, P. Gras, C. Combes, D. Laurencin, *Chem. Commun*. 2018, 54, 9591.

4 C. Gervais, D. Laurencin, A. Wong, F. Pourpoint, J. Labram, B. Woodward, A.P. Howes, K. Pike, R. Dupree, F. Mauri, C. Bonhomme, M.E. Smith, *Chem. Phys. Lett.* 2008, 464, 42.

