## Supplementary information for "Metabolic acids impact bone mineral maturation"

### Supplementary Information
#### for
### Metabolic acids impact bone mineral maturation


Yang Li[1], Rui Li[1], Karin H. Müller[2], Danielle Laurencin[3], Christian Bonhomme[4] and Melinda J Duer[1]*

[1]*Department of Chemistry, University of Cambridge, Lensfield Road, Cambridge, CB2 1EW, UK*
[2]*Cambridge Advanced Imaging Centre, Department of Physiology, Development and Neuroscience, University of Cambridge, Downing Street, Cambridge, CB2 3DY, UK*
[3]*ICGM, Univ Montpellier, CNRS, ENSCM, Montpellier, France*
[4] *Laboratoire de Chimie de la Matière Condensée de Paris, UMR Sorbonne Université CNRS, Sorbonne Université, 4, place Jussieu, 75252 Paris Cedex 05*, *France*


**Table S1:** Elemental and pXRD analyses of the OCP-metabolic acid materials in this work. Ca/P ratio, number of carbon and phosphorus atoms per unit cell are calculated from elemental analysis. $d_{100}$ lattice spacings for the 48-hour and 10-day synthesis materials are determined from powder XRD (Fig 2; see SI for details). Carbon, calcium and phosphate compositions are averages from typically three separate syntheses.

| Material | Carbon wt% 48-hour synthesis | Carbon wt% 10-day synthesis | Approx. no. of carbon atoms per unit cell§ 48-hour synthesis (10-day synthesis) | Ca/P ratio* 10-day synthesis | No of P / unit cell§ 10-day synthesis | $d_{100}$ / nm‡ 48-hour synthesis | $d_{100}$ / nm‡ 10-day synthesis |
|---|---|---|---|---|---|---|---|
| OCP-citrate | 3.5 | 3.6 | 6.2 (6.4) | 1.49 | 10.8 | 2.15 | 2.18 |
| OCP-lactate | 1.7 | 2.5 | 3.0 (4.4) | 1.41 | 11.4 | 1.92 | 1.87 |
| OCP-citrate-lactate | - | 3.4 | - (5.9) | 1.50 | 10.7 | - | - |

§ Per unit cell equivalents of the metabolic anion and P content is given for comparison to the original OCP structures, and so a unit cell is assumed to contain 16 $Ca^{2+}$ as in OCP.

* Ca/P ratio for pure OCP is 1.33; for OCP with substitution of one $HPO_4^{2-}$, the Ca/P ratio is 1.45; with substitution of two $HPO_4^{2-}$, Ca/P ratio is 1.6. For comparison, the Ca/P ratio for HAp is 1.67 and for the $\alpha$-tricalcium phosphate (TCP) starting material in the OCP-metabolic acid syntheses, 1.5.

‡ $d_{100}$ for pure OCP is 1.87 nm.

The carbon content of the 10-day synthesis with citrate is equivalent to approximately one citrate anion (6 carbons) per OCP unit cell, with lactate to approximately 1.5 lactate anions (3 carbons per lactate) per unit cell and for the mixed citrate-lactate synthesis, approximately one citrate or two lactate anions per unit cell, or some linear combination thereof. TGA showed typically 10 wt% water in the 10-day synthesis materials, but the water content measured by TGA is highly variable between samples. 10 wt % water corresponds to ~10 molecules of water per unit cell if the initial OCP-metabolic acid double salt chemical composition is assumed, and so broadly similar to that of the initial OCP-metabolic acid double salts.[1,2]

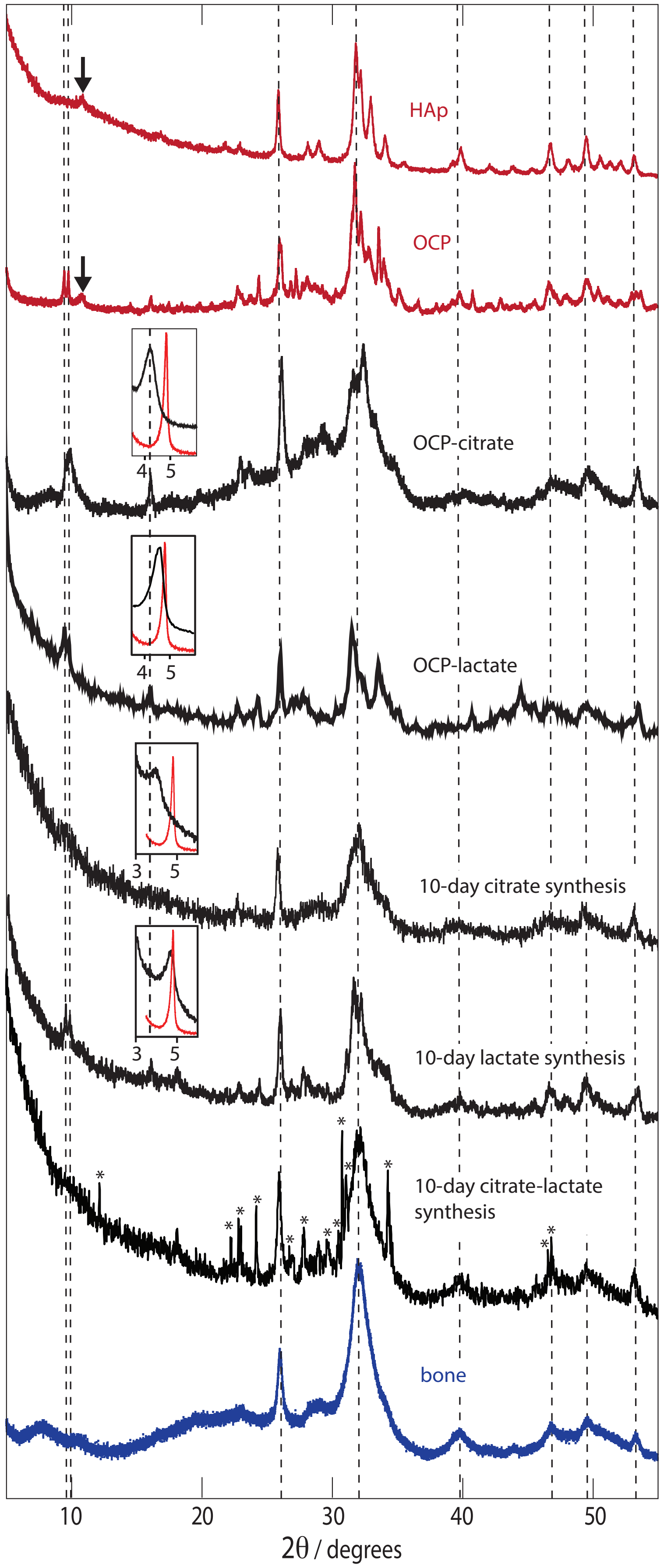
HAp
OCP
OCP-citrate
4
5
OCP-lactate
4
5
10-day citrate synthesis
3
5
10-day lactate synthesis
3
5
10-day citrate-lactate
synthesis
bone
10
20
30
40
50
2θ / degrees

**Fig S1:** pXRD diffraction patterns for the OCP-metabolic acid double salts (2-day syntheses) and the 10-day synthesis materials. Insets: the (low angle) (100) reflection (black) compared to that for pure OCP (red); horizontal scales are $2\theta$ in degrees. Dotted lines indicate the strongest reflections from bone (mineral) and two reflections around $2\theta$ ~10° that are distinctive for the OCP crystal structure ((1-10) and (010)). Arrows in the patterns for HAp and OCP indicate the putative HAp peak in the OCP diffraction pattern. The diffraction pattern for 10-day synthesis OCP-citrate-lactate has residual (sharp) reflections from $\alpha$-TCP; the expected positions of these are marked with *. The dotted line on the insets is to show how the $d_{100}$ reflection position changes between the *48 hour* and *10 day* syntheses for the citrate-containing materials.The pXRD patterns of both OCP-citrate and OCP-lactate double salts exhibit the characteristic OCP low angle (100) reflection between $2\theta$ ~ 4 - 5° ($2\theta$ = 4.839° for OCP; see Figure 2 insets) and strong OCP-like (1-10) and (010) reflections around $2\theta$ ~10° ($2\theta$ = 9.599°, 9.831° for OCP).

The $d_{100}$ spacings are larger in OCP-citrate than pure OCP (OCP $d_{100}$ = 1.87 nm; OCP-lactate $d_{100}$ = 1.92 nm; OCP-citrate $d_{100}$ = 2.15 nm; see Table 1) consistent with incorporation of the associated metabolic acid anion into the OCP hydrated layer. The $d_{100}$ spacing for OCP-citrate is larger than that for OCP-lactate consistent with citrate being a larger anion than lactate. Samples of pure OCP typically contain some small amount of HAp.[3,4] That is the case here too, evidenced, by a small, broad HAp (100) reflection in the pXRD pattern for the OCP sample at $2\theta$ ~10.9° (other expected HAp reflections overlap with the relatively broad OCP reflections).

For the 10-day lactate-containing synthesis, the pXRD pattern contains the distinctive OCP-like (100) (see inset in Fig S2), (1-10) and (010) reflections (see Fig S2). For this 10-day synthesis material, the $d_{100}$ spacing is smaller than in the initial OCP-lactate double salt. The pXRD reflections for the 10-day synthesis lactate material are generally broadened compared to the reflections for the OCP-lactate double salt, particularly in the $2\theta$ ~ 30° - 35° range where some of the more distinctive reflections that would distinguish an HAp or OCP-like structure are expected. Above $2\theta$ ~ 30°, the differences in the pXRD patterns for pure HAp and OCP are indeed subtle. This along with the broadening of reflections means that between $2\theta$ ~ 30° - 60°, the 10-day synthesis lactate material pXRD pattern resembles both the HAp and OCP pXRD patterns; only the distinctive OCP-like (100), (1-10) and (010) reflections at low $2\theta$ suggest predominant retention of OCP structural characteristics. For the 10-day synthesis citrate material, a distinctive, though broad, OCP-like (100) reflection remains at low angle. The associated average $d_{100}$ spacing is slightly larger than for the OCP-citrate double salt (2.18 nm compared to 2.15 nm in the OCP-citrate double salt, Table 1). There are no other reflections that can be confidently assigned to specifically OCP-like structures or HAp. The reflections are broad so that as in the case for the 10-day synthesis lactate materials, the pXRD pattern for the 10-day citrate material resembles those for both HAp and OCP above $2\theta$ ~ 30°. The 10-day mixed citrate-lactate-containing synthesis product did not show any low-angle reflections consistent with OCP-like (100), (1-10) and (010) reflections and the pXRD pattern for this material overall resembles more that for disordered/ nanocrystalline HAp (Fig 2) than an OCP-like material. Some weak reflections from the $\alpha$-TCP starting material are often present in this diffraction pattern, indicating that there has been incomplete reaction of the $\alpha$-TCP.

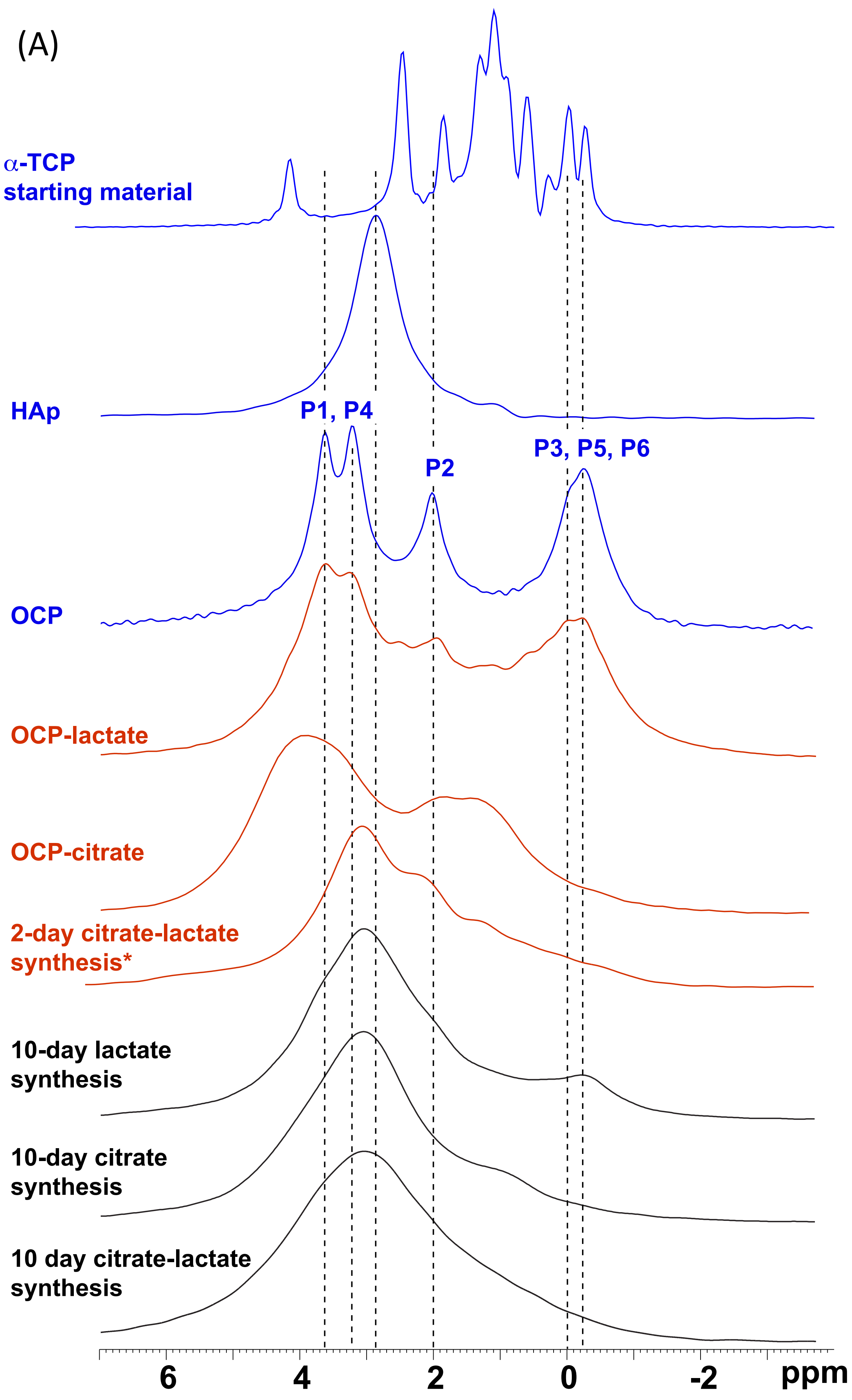
(A)
α-TCP
starting material
HAp
P1, P4
P2
P3, P5, P6
OCP
OCP-lactate
OCP-citrate
2-day citrate-lactate
synthesis*
10-day lactate
synthesis
10-day citrate
synthesis
10 day citrate-lactate
synthesis
6
4
2
0
-2
ppm

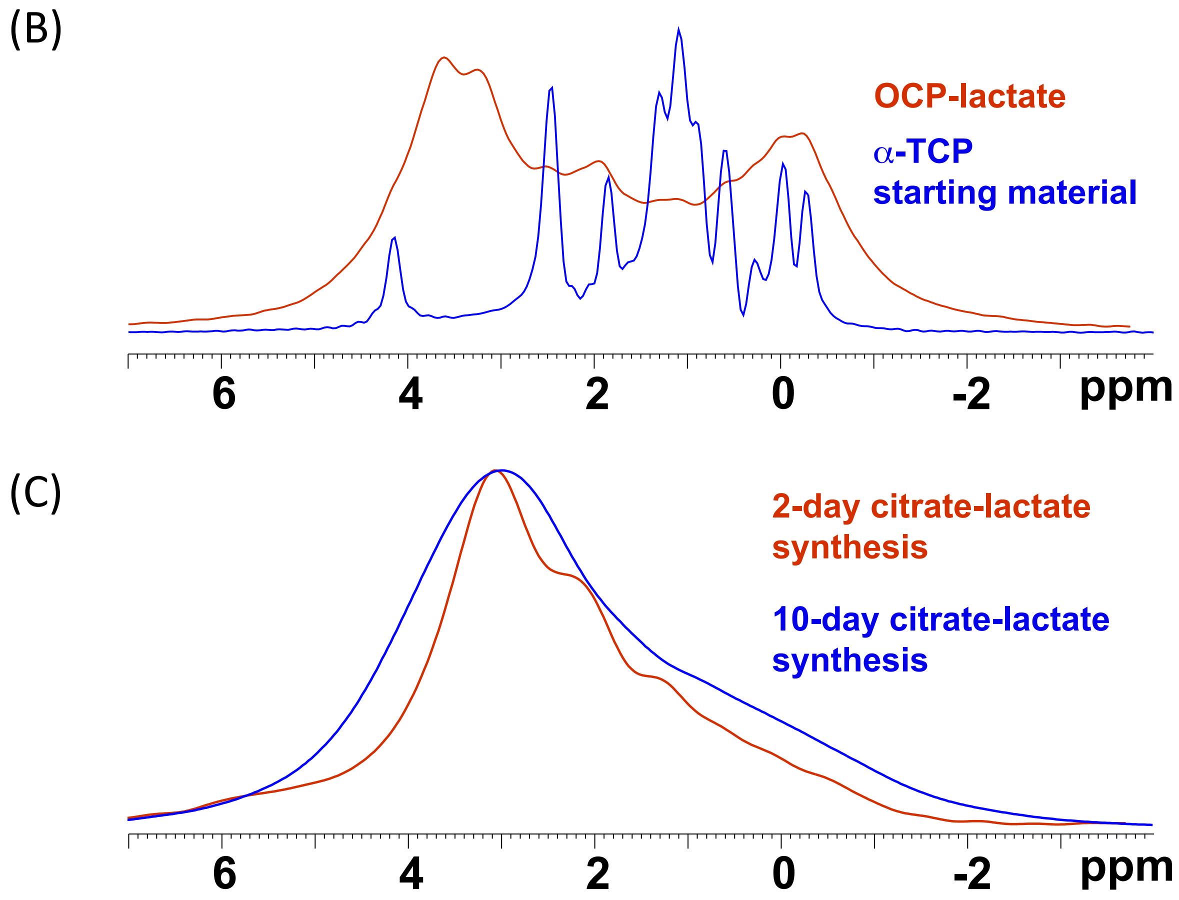


**Figure S2: (A)** Comparison of the $^{31}P$ DP MAS NMR spectra of the OCP-metabolic acid double salts and the 10-day synthesis materials investigated in this work. * $^{31}P$ CP MAS NMR spectrum; 48-hour synthesis containing a mixture of citrate and lactate led to only very small amounts of product with the α-TCP reactant being the only compound observed by DP $^{31}P$ NMR, hence the $^{31}P$ CP NMR spectrum is shown here, because signals from α-TCP are not observed by cross polarization. Sample spinning rate is 10 kHz for all samples. $^{31}P$ DP spectra for pure HAp (nanocrystalline) and OCP are shown for reference at the top. Dotted lines indicate the chemical shifts of the characteristic HAp and OCP $^{31}P$ signals. Note that the spectrum for the OCP-lactate double salt after 2 days shows residual signals from the α-TCP starting material used in the synthesis (see Fig S2B for overlay of the OCP-lactate and α-TCP $^{31}P$ spectra ). OCP assignments refer to the phosphate labelling in Fig 1. (B) Overlay of the $^{31}P$ direct polarization spectra for the 2-day synthesis OCP-lactate sample and the a-TCP starting material, showing that there are residual signals from the starting material present in the OCP-lactate sample (see Fig 4, main text). (C) Overlay of the CP spectra for the 2 and 10-day synthesis citrate-lactate sample for comparison (at the 2-day time point, there is still a large amount of the a-TCP starting material remaining making it difficult to detect the OCP-lactate product with direct polarization at this synthesis time point; CP precludes signals from a-TCP because a-TCP contains no $^{1}H$).
Fig S2:

The $^{31}P$ DP MAS spectrum for the OCP-citrate double salt has been reported and assigned previously:[1] in summary, the highest frequency $^{31}P$ chemical shifts (signal/s in the range 3.1 – 3.6 ppm) are from the OCP apatitic orthophosphate groups (P1, P4, see Fig 1 for phosphate labelling), the lowest frequency signals (broad, poorly resolved set of signals, 0 - –1 ppm) are from the hydrated layer $HPO_4^{2-}$ / $PO_4^{3-}$ (P3 ($PO_4^{3-}$); P5, P6 ($HPO_4^{2-}$)) and the intermediate frequency signal (1.8 ppm for OCP-citrate, 2.0 ppm in pure OCP) is from the orthophosphate groups in the interface between the OCP apatitic-like and hydrated layers (P2). The OCP-lactate double salt/ 48-hour synthesis product gives a similar distribution of $^{31}P$ chemical shifts, and we assign them similarly (note that there are some small residual signals from α-TCP also in the $^{31}P$ DP NMR spectrum of the OCP-lactate double salt/ 48-hour synthesis sample (Fig 4 and S2A)). The 48-hour synthesis with both citrate and lactate in the reaction mixture led to only small amounts of product as described above so that $^{31}P$ DP spectra were dominated by the signals from the solid α-TCP starting material. Thus, we instead recorded $^{31}P$ cross-polarization (CP) NMR spectra of the products of 48-hour synthesis with the citrate-lactate mixture as CP prevents signals from α-TCP being observed (there are no $^{1}H$ in α-TCP so $^{1}H\{^{31}P\}$ CP cannot generate signals from α-TCP). The peak maximum in the orthophosphate region (3.05 ppm) of the resulting CP $^{31}P$ spectrum is between that expected for OCP-like (3.1 – 3.6 ppm) and HAp (2.85 ppm) orthophosphate sites, and is similar to that expected for nanocrystalline HAp.[5–9] Broad signals centred at ~2.2 and ~1.3 ppm, plus the low frequency "tail" in the spectrum cover the spectral region for OCP-like hydrated layer $HPO_4^{2-}$ and $PO_4^{3-}$ and hydrated layer-apatitic layer interface region, suggesting there are OCP-like structures in the material. The CP relative intensities are not quantitative so we cannot deduce the relative amounts of the different OCP phosphatic environments from this spectral analysis.

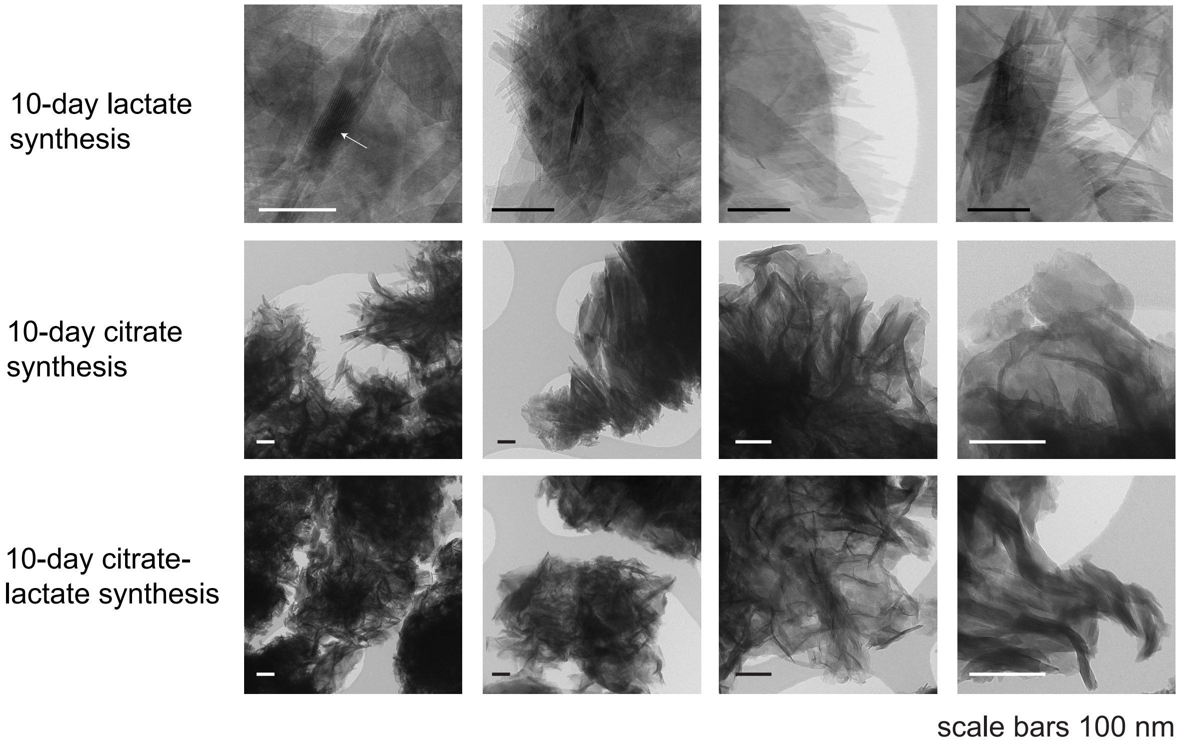


**Figure S3: Additional TEM images of 10-day synthesis materials**. For the lactate-only material the white arrow indicates a particle that exhibits interlayering. All scale bars, 100 nm. To note, any remaining α-TCP crystals in the mixed citrate-lactate samples are expected to be much larger than the lengthscales represented by the sizes of the TEM images; the pXRD reflections from α-TCP crystals/ domains in these samples are very sharp (Fig S1) corresponding to crystals with sizes much larger than the few hundred nanometer image dimensions here.

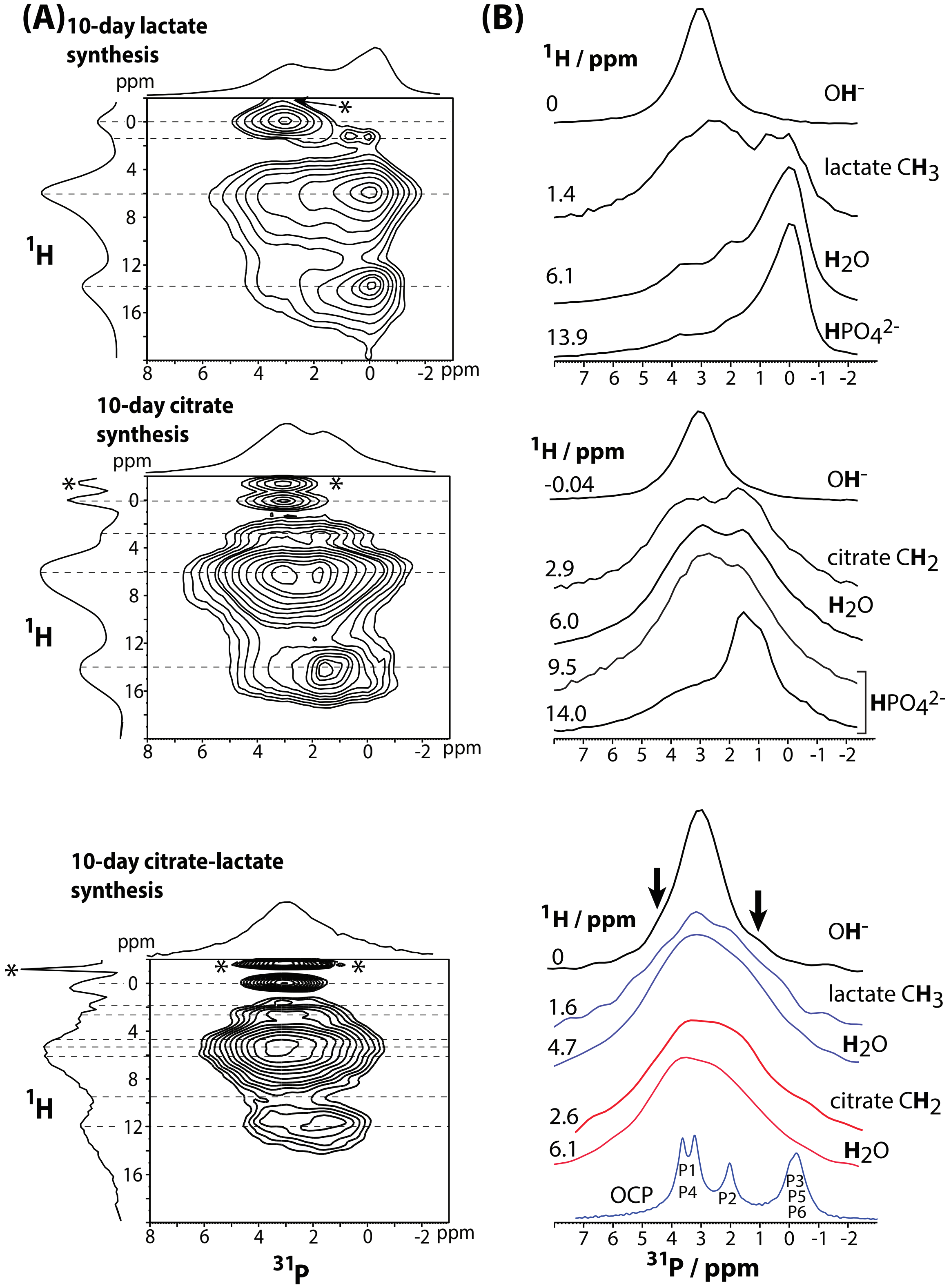


**Fig S4:** 2D $^1$H-$^{31}$P correlation spectra for the 10-day materials synthesized in this work. The 2D $^1$H-$^{31}$P correlation spectrum for the 10-day synthesis lactate material shows a $^1$H 0 ppm – $^{31}$P 3.1 ppm correlation consistent with HAp structures, $^1$H 0 ppm corresponding to the HAp $OH^-$ $^1$H signal and $^{31}$P 3.1 ppm to HAp orthophosphate in nanocrystalline material. [5–9] The broadness of the HAp $^{31}$P signal compared to that typical of crystalline HAp (e.g. see Fig S2), indicates that the HAp orthophosphate sites are disordered or heterogeneous, as would be expected for nanoscopic-sized domains. There is also an unequivocal $^1$H signal from $HPO_4^{2-}$ centred at 13.9

ppm. This $^1H$ signal is correlated with $^{31}P$ signals (3.5, 2.0, 0.5 ppm), that may be assigned to OCP-lactate-like environments (see the 48-hour lactate only synthesis spectrum in Fig S2). Signal intensity in these spectral slices depend on the spatial proximity of the $^{31}P$ sites to the particular $^1H$ site as well as the abundance of the $^{31}P$ site; the closer the $^1H$ and correlated $^{31}P$ are in space, the higher the correlation spectral intensity because the correlation intensity derived from CP from $^1H$ to $^{31}P$. The lower frequency $^{31}P$ signals in this correlation are from $HPO_4^{2-}$ and $PO_4^{3-}$ strongly hydrogen bonded to water $^1H$ (0.5 – 2.0 ppm) and have the highest intensity in the correlation by virtue of these $^{31}P$ sites being spatially closest to $HPO_4^{2-}$ $^1H$ in the OCP-lactate structure. The water $^1H$ signal at 6.1 ppm is consistent with water in an OCP-like hydrated layer and correlates with a similar intensity distribution of $^{31}P$ signals as the $HPO_4^{2-}$ $^1H$ signal, showing that the water is in a similar environment to the $HPO_4^{2-}$, namely an OCP-like hydrated layer. The lactate methyl $^1H$ signal at 1.4 ppm shows correlation with a very broad/ poorly resolved set of $^{31}P$ signals covering the expected range for HAp- and OCP-like structures. Importantly, there is a much stronger correlation with the $^{31}P$ chemical shift range for apatitic orthophosphate (2.8 – 3.6 ppm) than would be expected if the lactate were confined to an OCP-like hydrated layer; if it were, we would expect an intensity distribution in the spectral slice more similar to those for the $HPO_4^{2-}$ and water $^1H$ spectral slices. We suggest that at least some lactate is at interfaces with HAp, resulting in relatively strong spectral correlation between lactate $^1H$ and HAp-like orthophosphate $^{31}P$. We also note that structured water $^1H$ on an HAp surface can give a signal at ~1.4 ppm,[7,13] though the amount of HAp in this sample would suggest that there would be relatively few such sites compared to the amount of lactate (approximately one lactate per unit cell) and therefore that such a signal would be relatively low in intensity. Thus, we conclude that the product of the 10-day lactate-only synthesis is dominated by molecular structures similar to the initial OCP-metabolic acid double salts with some additional HAp-like regions, consistent with the pXRD, TEM and 1D $^{13}C$, $^{31}P$ and REDOR NMR results discussed above, with lactate probably present near HAp domains.

The 2D $^1H$-$^{31}P$ correlation spectrum for the 10-day synthesis citrate material shows a similar $^1H$ (0 ppm) – $^{31}P$ (3.1 ppm) correlation signal as for the 10-day synthesis lactate material, consistent with nanocrystalline hydroxyapatite structures. The $^{31}P$ lineshape in this correlation is broad as for the 10-day lactate-only material, consistent with nanoscopic-sized HAp domains which TEM confirms exist in this material (Fig 2). The broad $^1H$ signal centred at 14.2 ppm is unequivocally due to $^1H$ in OCP-like $HPO_4^{2-}$ anions, confirmed by the most intense $^{31}P$ signals correlated with them being between 0.9 – 1.5 ppm, the expected chemical shift range for OCP-like hydrated layer $HPO_4^{2-}$ and $PO_4^{3-}$ (P3, P5, P6 in Fig 1) as for the lactate material. There are also the expected weaker correlations to $^{31}P$ signals above 3 ppm consistent with OCP apatitic-like $PO_4^{3-}$ (P1, P4 in Fig 1) that are more distant from the hydrated layer $HPO_4^{2-}$ $^1H$, and therefore exhibit weaker spectral correlations to hydrated layer $^1H$. There is $^1H$ signal intensity in the range 9 – 10 ppm consistent with $HPO_4^{2-}$ $^1H$ on the surface of nanocrystalline HAp.[7,13] $^1H$ signals in this chemical shift range correlate with a broad range of $^{31}P$ chemical shifts and this is discussed in more detail below in the context of the mixed citrate-lactate material where the equivalent signal is more intense. The citrate $CH_2$ $^1H$ signal at 2.9 ppm[1] is correlated with a set of broad overlapping $^{31}P$ signals covering the chemical shift range for OCP-citrate and HAp structures, consistent with citrate being both in OCP-citrate-like hydrated layers and at internal OCP hydrated layer interfaces with HAp as for the lactate material. The $^1H$ signal for water is centred at 6.0 ppm, but broad and asymmetric, suggesting multiple water environments. A $^1H$ chemical shift of 6 ppm is consistent with the water $^1H$ signal for the hydrated layers of OCP-citrate, some of which undoubtedly remain in this material from pXRD (Fig S1) and 1D $^{31}P$ NMR (Fig S2).[1] By way of comparison, water molecules on external surfaces of nanocrystalline HAp have a $^1H$ chemical shift of 4.85 ppm.[14] In contrast to the 10-day synthesis lactate material however, this 6.0 ppm water $^1H$ signal correlates strongly with the $^{31}P$ chemical shift range for apatitic orthophosphate (2.8 – 3.6 ppm) as well as the expected $^{31}P$ signals from the OCP hydrated layer phosphatic sites. As OCP apatitic orthophosphate sites are relatively distant from the OCP water, there is not expected to be a strong spectral correlation between OCP water $^1$H and OCP apatitic $^{31}$P signals, as indeed we found for the 10-day synthesis lactate material (and see for example the NMR data for the OCP citrate double salt in reference [1]). We thus suggest that this strong spectral correlation between OCP-like water and apatitic orthophosphate $^{31}$P must arise from OCP-like water, i.e. strongly hydrogen bonded water/ trapped water, that is close in space to HAp orthophosphate. This assignment is consistent with the interlayered crystal morphologies seen by TEM (Fig 2) for the 10-day synthesis citrate material, and suggests that in the interlayered structure, it is the OCP hydrated layers that predominantly interface with the HAp domains rather than the OCP apatitic layers. We conclude that whilst there are still identifiable OCP-like structures present in the 10-day synthesis citrate material, there is a significant amount of HAp structure which interfaces with OCP-like hydrated layers and citrate.

The 2D $^1$H-$^{31}$P correlation spectrum for the 10-day synthesis mixed citrate-lactate material again has a characteristic $^1$H 0 ppm – $^{31}$P 3.1 ppm correlation signal consistent with a nanocrystalline HAp component. Importantly, there is also an observable correlation between the HAp $OH^-$ $^1$H signal and a broad $^{31}$P shoulder at ~ 1.2 ppm, a $^{31}$P chemical shift similar to those for the hydrated orthophosphate and $HPO_4^{2-}$ in OCP-like hydrated layers in the citrate-only material (B, 0 ppm $^1$H slice, arrow) consistent with HAp domains interfacing with OCP-like hydrated layers as for the citrate-only material. There is $^1$H signal characteristic of $HPO_4^{2-}$ from ~9 – 14 ppm, suggesting multiple $HPO_4^{2-}$ populations, as for the citrate-only 10-day material. The lower frequency $HPO_4^{2-}$ $^1$H signal range (9 – 11 ppm; Fig S3B) consistent with $HPO_4^{2-}$ on HAp surfaces,[7,13] correlates with a broad $^{31}$P signal centred at ~ 3 ppm, characteristic of (disordered) HAp-like orthophosphate and with significant spectral intensity to higher chemical shift including a putative broad shoulder at ~5.5 ppm (B, 9.5 ppm $^1$H slice, arrow) which is the expected $^{31}$P chemical shift range for $HPO_4^{2-}$ on HAp surfaces.[7,13] The higher $^1$H chemical shifts in the $HPO_4^{2-}$ signal region (12 - 14 ppm) are consistent with OCP-like hydrated layer $HPO_4^{2-}$ $^1$H (as for the citrate-only and lactate-only materials) but correlate with a set of broad poorly resolved $^{31}$P signals centred at: ~3.7 ppm, ~2.7 ppm and ~1.4 ppm. The $^{31}$P 1.4 ppm signal is consistent with OCP-like hydrated layer orthophosphate. The 3.7 and 2.7 ppm signals are consistent with apatitic-like orthophosphate of some sort and/ or $HPO_4^{2-}$ substitutions in HAp (expected to be ~2.5 – 5.5 ppm).[7,13] Importantly, the $^1$H 12 – 14 ppm signal range correlates strongly with all of the 3.7, 2.7 and 1.4 ppm $^{31}$P signals. If the $^{31}$P signals in the 2.7 – 3.7 ppm region were from OCP-like apatitic orthophosphate only, we would expect to see relatively weak correlations to them from OCP hydrated layer $HPO_4^{2-}$ $^1$H, as for the citrate- and lactate-only materials. Thus, the 2.7 and 3.7 ppm-centred $^{31}$P signals must be from sites closer to the remaining OCP hydrated layers than OCP-like orthophosphate and likely more prevalent than in the citrate-only and lactate-only materials, to account for the strong spectral correlation from OCP hydrated layer $HPO_4^{2-}$ $^1$H. We thus assign the $^{31}$P signals centred at ~2.7 and ~3.7 ppm to HAp orthophosphate and $HPO_4^{2-}$ near the interface with OCP hydrated layers in the interlayered structure of this mixed citrate-lactate material. $^{31}$P chemical shifts around 3.7 ppm are consistent with $HPO_4^{2-}$ substitutions in HAp which could be prevalent given the formation of HAp here from an initial $HPO_4^{2-}$-containing OCP lattice. $^{31}$P chemical shifts around 2.7 ppm are intermediate between that for nanocrystalline HAp orthophosphate (3.1 ppm) and the OCP P2 orthophosphate site (2 ppm in pure OCP, 1.8 ppm in OCP-citrate and 1.9 ppm in OCP-lactate 48-hour double salts). Thus, $^{31}$P signals around 2.7 ppm are consistent with orthophosphate sites with environments intermediate between those in HAp and OCP, i.e. orthophosphate less strongly hydrogen bonded than in the hydrated orthophosphate OCP P2 site/ more strongly hydrogen bonded than bulk HAp orthophosphate, which would be consistent with HAp orthophosphate at HAp-OCP interfaces. The proposed assignments for the phosphatic sites for the mixed citrate-lactate 10-day synthesis material and their approximate spatial distributions are depicted schematically in Fig S5.

Importantly, the water $^1$H signal for the 10-day mixed citrate-lactate material is centred at 5.4 ppm, which is a significantly lower chemical shift than the ~6 ppm water $^1$H chemical shift for OCP-citrate and OCP-lactate. However, the water $^1$H signal is notably broader than for the citrate-only and lactate-only materials, and somewhat asymmetric, with significant intensity to higher chemical shift as well. The lower water $^{1}H$ chemical shifts are similar to the (external) surface water $^{1}H$ chemical shifts for nanocrystalline HAp (~4.85 ppm)[6] and correlate with $^{31}P$ signals centred ~2.9 ppm suggestive of HAp-like orthophosphate, e.g. $^{1}H$ 4.7 ppm in (B) for this material; thus we assign the lower range of water chemical shifts (below ~5 ppm) to external surface water on HAp domains. Higher water $^{1}H$ chemical shifts, e.g. 6.1 ppm in (B) for this material, correlate with a broad range of signals between ~0 ppm to over 5 ppm, but most strongly $^{31}P$ chemical shifts between ~2.5 and ~3.7 ppm, similar to the higher frequency $^{31}P$ signals to which the $^{1}H$ 12 – 14 ppm $^{1}H$ signal from OCP hydrated layer $HPO_4^{2-}$ correlates. This is consistent with the 6.1 ppm water $^{1}H$ correlation here being between OCP-like hydrated layer water $^{1}H$ and HAp phosphatic sites near HAp-OCP interfaces as described above for the $^{1}H$ 12 – 14 ppm correlation.

The citrate $CH_2$ and lactate methyl $^{1}H$ signals (~2.6 and ~1.6 ppm respectively) overlap with the expected $^{1}H$ chemical shifts for HAp structured surface water ($H_2O$ substituting for $OH^-$ and HAp surfaces) and $HPO_4^{2-}$ substitutions in HAp, as well as some overlap with the broad water $^{1}H$ signals for this material. Thus, we cannot conclude what part of the correlated $^{31}P$ spectra are from citrate and lactate $^{1}H$ or mineral $^{1}H$. Nevertheless, that there is significant 1H signal intensity at the $^{1}H$ chemical shifts expected for citrate and lactate $^{1}H$ demonstrates that both are present in the mineral structure.

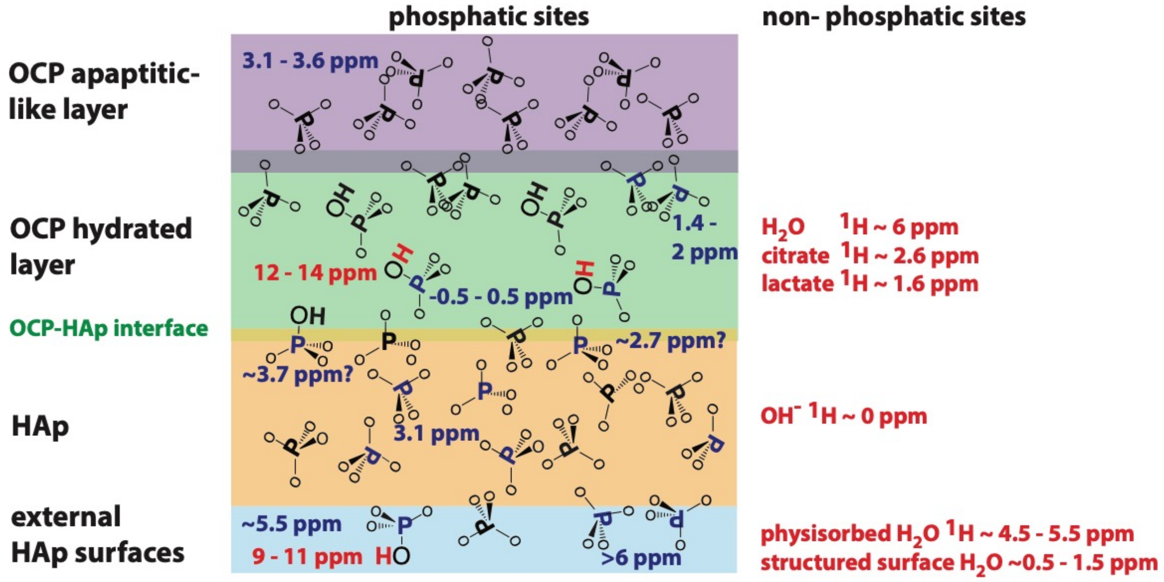


**Fig S5:** Schematic depiction of the proposed $^{31}P$ and $^{1}H$ NMR assignments for the mixed citrate-lactate 10-day synthesis material. $^{31}P$ chemical shifts are in blue, $^{1}H$ in red. Note that figure is not intended to illustrate the morphologies of the OCP or HAp regions. Which HAp crystal surfaces are external surfaces depends on how the OCP and HAp regions are interfaced. HAp structured surface water molecules are those that substitute for $OH^-$ groups at HAp surfaces.

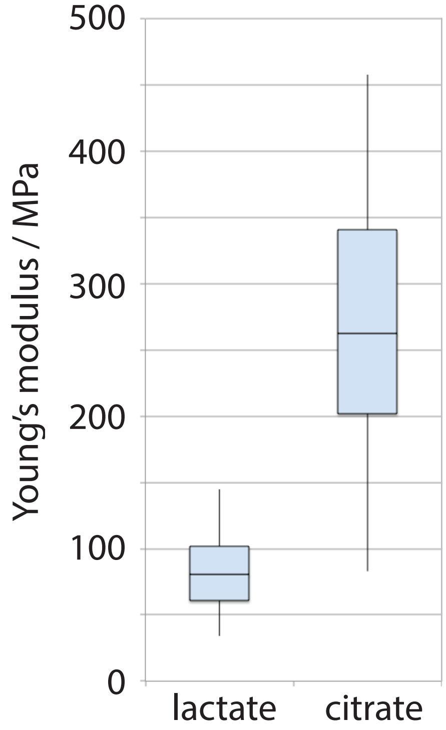


**Figure S6:** AFM-derived stiffness modulus for the 10-day synthesis citrate-only and lactate-only materials.
